# REGULATORY T CELLS PROTECT AGAINST ABERRANT REMODELING IN A MOUSE MODEL OF PULMONARY FIBROSIS

**DOI:** 10.1101/2025.07.02.662777

**Authors:** Aditi Murthy, Luis R. Rodríguez, Willy Roque Barboza, Yaniv Tomer, Sarah Bui, Paige Carson, Thalia Dimopoulos, Swati Iyer, Katrina Chavez, Charlotte H. Cooper, Jeremy B. Katzen, Michael F. Beers

## Abstract

Regulatory T (Treg) cells are well recognized for their role in immune regulation; however, their role in tissue regeneration is not fully understood. This study demonstrates such a role of Tregs in a published preclinical murine model of spontaneous pulmonary fibrosis (PF) expressing a human PF related mutation in the Surfactant Protein-C (SP-C) gene (*SFTPC^I73T^*). Genetic crosses of SP-C^I73T^ mice with Foxp3^GFP^ and Foxp3^DTR^ lines were utilized to study Treg behavior during PF development. We found that FoxP3+Tregs accumulate during the transition from inflammation to fibrogenesis, peaking at 21-28 days after mutant *Sftpc^I73T^* induction localizing to both perivascular and distal fibrotic lung regions. Diphtheria toxin mediated ablation of Tregs at 17 days worsened fibrosis and increased levels of TGFβ and inflammatory cytokines. Tregs expressed Th2 markers (Gata3+) and elaborated factors including amphiregulin (Areg) and Osteopontin (Spp1). Reductionist experiments showed that lung Tregs enhanced organoid formation when co-cultured with alveolar epithelial cells and adventitial fibroblasts, an effect size mimicked using Areg and Spp1 in combination. Our findings demonstrate that immune-mesenchymal-epithelial signaling crosstalk is present in the distal lung wherein Tregs play a protective role by limiting fibrosis and promoting tissue repair, highlighting their broader function beyond immune modulation in lung injury.

**HIGHLIGHTS:**

- In a preclinical model of spontaneous pulmonary fibrosis, regulatory T cells (Tregs) were found to infiltrate the lung coincident with the resolution of early injury and transition to fibrogenesis.
- Depletion of Tregs at this transition worsened lung injury and enhanced fibrogenesis.
- Tregs recovered from the fibrotic lung are Type 2 skewed – GATA3+ and produce growth factors (e.g. Amphiregulin, Osteopontin) that promote lung tissue repair in ex vivo organoid models.

## INTRODUCTION

Aberrant fibrotic remodeling is a frequent complication of and major contributor to the morbidity and mortality of a number of chronic interstitial lung diseases (ILDs) in the aging adult lung [1, 2]. Among these, the most pernicious is Idiopathic Pulmonary Fibrosis (IPF) defined by a histopathology containing a characteristic disruption of distal lung architecture and a distinct high resolution CT radiographic pattern that ultimately leads to abnormal gas exchange, respiratory failure, death and/or need for transplantation [3]. These devastating outcomes have been remarkably refractory when current “gold-standard” therapy based on “antifibrotic” strategies is employed [4–6]. The dearth of IPF therapeutics can be linked in part to an incomplete understanding of its pathophysiologic underpinnings. While it is now appreciated that IPF represents a dysfunctional interplay between epithelial, mesenchymal, and effector cell lineages in the distal lung to generate a “fibrotic niche“, the most recent emphasis on epithelial-mesenchymal crosstalk has left open questions regarding the roll of immune cell populations in the cascade of events that produce end-stage lung scarring.

Previous work has indicated a role for myeloid lineages in lung homeostasis, injury, and aberrant repair [7, 8]. However, the contributions of lymphoid populations to reparative or fibrotic lung niche is less defined. Regulatory T cells (Tregs) are a distinct population of CD4+ lymphocytes which express the transcription factor forkhead homeoboxprotein-3 (Foxp3), are essential for maintaining immune homeostasis, and contribute to the prevention of autoimmunity [9]. While traditionally studied and identified for their role in immunosuppression and in the modulation of immune responses to prevent tissue damage, recent studies have highlighted their role in tissue repair following injury across various organs, including cardiovascular ischemia [10, 11], UV-induced skin injury [12], toxin-induced muscle injury [13], and neurological diseases [14, 15]. In these models, Tregs were found to interact with structural cells (i.e. epithelial, mesenchymal, and/or endothelial lineages) underscoring their context-dependent role in tissue homeostasis [16].

Similar observations are emerging for Tregs in the lung, but the data remains incomplete. In acute lung injury models, such as that induced by LPS, lymphocytes play a critical role in recovery, with Rag-/- mice exhibiting impaired recovery, increased mortality, weight loss, and prolonged histopathological damage [17]. Adoptive transfer of splenic CD4+CD25+ T cells rescues LPS-challenged mice [17]. This same group also showed that CD103+ Tregs regulate alveolar epithelial cell proliferation post-LPS injury [18] and that Treg-specific Fibroblast Growth Factor 7 (FGF-7;KGF) expression is vital for epithelial proliferation during injury resolution [19]. In a flu injury model, Treg-generated Amphiregulin (Areg), an EGFR ligand, was found to be crucial for maintaining lung function and preserving tissue morphology via an IL-18 and IL-33 signaling hub capable of driving Col14a+ (adventitial) fibroblasts to support alveolar epithelial organoid growth *in vitro* [20, 21]. Collectively, the data support a role for Tregs as mediators of tissue repair after injury through interactions with structural parenchymal cells to restore the alveolar niche. While Tregs are present in the human IPF lung [22], their role in its pathogenesis remains less well defined due in part to both challenges with traditional preclinical models (bleomycin injured rodents) and limitations associated with usage of end stage human IPF tissue or employment of ex vivo models (e.g. precision cut lung slices (PCLS) or lung organoids) largely devoid of relevant immune cell populations. To better understand the role for Tregs in lung fibrosis while also circumventing confounding technical issues, we leveraged previously developed inducible mouse models expressing high effect size variants in the surfactant protein C (*SFTPC*) gene found in patients with familial pulmonary fibrosis (FPF), sporadic IPF, and childhood interstitial lung disease (chILD), the most common of which is a missense variant in the linker domain of the surfactant protein C (SP-C) pro-protein (*SftpcI73T*)[23–25]. The *Sftpc^I73T^* model which is driven by intrinsic AT2 cell dysfunction stands in contrast to prior foundational models of lung fibrosis that require exogenous lung injury such as bleomycin, radiation, or silica. Mutant Sftpc^I73T^ mice have been shown to exhibit a phenotype marked by an early multiphasic injury/alveolitis (7-14 days) followed by a transition to fibrosis (14-42 days).

In this study, we first defined the ontogeny of Treg dynamics during an AT2 epithelial cell intrinsic aberrant lung injury/repair sequence using flow cytometry to interrogate both *Sftpc^I73T^* mice as well as a novel *Sftpc^I73T^* / Foxp3-GFP genetic model. Marked increases in lung Tregs occurred 3-4 weeks after induction of mutant *Sftpc^I73T^*in AT2 cells. Subsequent ablation of Tregs using a Diptheria toxin-based approach initiated during the transition from inflammation to fibrogenesis worsened the disease phenotype consistent with a protective role. Further characterization of bulk isolated Tregs revealed them to be activated, type 2-skewed, and expressing transcripts for growth factors including Areg and SPP1. Further equipoise for a protective role for this Treg population in alveolar repair and regeneration was then obtained in a reductionist, *ex vivo* organoid model which supported immune-epithelial-mesenchymal crosstalk within the alveolar niche mediated by a number of key reparative signaling nodes. Collectively, this study has mechanistically defined a critical role for Foxp3 CD4 regulatory T cells in modulating the cellular events associated with the development of pulmonary fibrosis.

## MATERIALS AND METHODS

### In Vivo Murine Fibrosis Models

Tamoxifen (Tam) inducible SP-C^I73T^ fibrosis mice (I^ER^-*Sftpc^I73T^*) were generated as previously published [26]. Foxp3 GFP reporter mice (Foxp3^eGFP^) expressing green fluorescent protein (GFP) downstream of endogenous Foxp3 stop codon resulting in fluorescence tagging of all Tregs [27] were a gift from Taku Kambayashi (University of Pennsylvania) (Strain^#^ 006772, The Jackson Laboratory Bar Harbor ME) and crossed with I^ER^-*Sftpc^I73T^* to produce SP-C^I73T^-Foxp3^eGFP^ mice with homozygosity of both alleles.

To generate an I^ER^-SP-C^I73T^-Foxp3^DTR^ knock-in mouse line, we used B6.129(Cg)-*Foxp3^tm3(Hbegf/GFP)Ayr^*/J purchased from The Jackson Laboratory (strain^#^ 016958). *Foxp3^DTR^* knock-in mice express the human diphtheria toxin receptor (designated heparin-binding EGF-like growth factor [HBEGF]) and EGFP genes inserted into the 3’ untranslated region of *Foxp3* locus-without disrupting expression of the endogenous *Foxp3* gene [28]. These were crossed into the parent I^ER^-*Sftpc^I73T^* line to reach homozygosity of all alleles. Intraperitoneal administration of diphtheria toxin (DT) was used for specific depletion of Foxp3+ Tregs.

Male and female mice were included in all protocols. Tamoxifen treatment of I^ER^-*Sftpc^I73T^*, SP-C^I73T^- Foxp3^eGFP^ and SP-C^I73T^-Foxp3^DTR^ mouse lines was initiated at 12-14 weeks of age by oral gavage performed on days 0 and 4 as published [29]. All mice were housed under pathogen free conditions in an AALAC approved barrier facility at the Perelman School of Medicine, University of Pennsylvania. All studies were approved by the Institutional Animal Care and Use Committee at the University of Pennsylvania.

To generate cell populations for organoid cell culture assays, AT2 cells were obtained from bi-transgenic SP-C^CreERT2^ (Strain^#^ 028054) - Rosa26^tdT^ (Strain^#^ 007914, The Jackson Laboratory) mice as described below. Bulk fibroblast populations were harvested from PDGRFα^EGFP^ mice (Strain^#^ 007669, The Jackson Laboratory) as described below.

### Reagents and Materials

Antibodies used for flow-cytometry, FACS, and immunohistochemistry were obtained from commercial sources (**Supplemental Tables 1 and 3)**. Tamoxifen (non-pharmaceutical grade) was purchased from Sigma-Aldrich, Inc. (St. Louis MO). Diphtheria toxin (Catalog^#^ D0564) was purchased from MilliporeSigma (Burlington MA), reconstituted, and stored as per manufacturer’s instructions. Except where noted, all other reagents were electrophoretic or immunological grade and purchased from commercial sources as noted.

### Multichannel Flow Cytometry and Cell Sorting

Flow cytometry was performed as we described [26, 29, 30]. Blood free perfused lungs were digested in Phosphate Buffered Saline (Mg and Ca free) with 2 mg/ml Collagenase Type I (Gibco, Grand Island, NY) and 50 units of DNase (MilliporeSigma, Burlington MA), passed through 70-μm nylon mesh to obtain single-cell suspensions, and then processed with ACK Lysis Buffer (Thermo Fisher Scientific Waltham, MA). Cell pellets collected by centrifugation were resuspended in PBS+0.1% Bovine Serum Albumin (Jackson ImmunoResearch Laboratories Inc, West Grove, PA) and aliquots removed for cell counting using a NucleoCounter (New Brunswick Scientific, Edison, NJ). Cell pellets were incubated with anti-mouse CD16/CD32 Ab (Fc block) (eBioscience San Diego, CA) for 15 minutes at 4°C to block nonspecific binding. This was followed by live/dead staining. Cells were incubated for 30 minutes in dark with viability dye followed by incubation with antibody mixtures for cell surface antigens, 30 minutes at 4°C (or isotype controls) (see **Supplemental Table 1)**. Cells were fixed with 2% paraformaldehyde for 15 minutes at 4°C and then re-suspended in PBS+1%BSA+0.1% sodium azide (Sigma-Aldrich) buffer before analysis on a LSR Fortessa (BD Biociences, Franklin Lakes, NJ). For intracellular staining cells were fixed and permeabilized with a Foxp3 buffer set (eBiosciences) following manufactures protocol. FMO controls were used for intracellular markers. Dynabeads Untouched Mouse T cell kit (ThermoFisher Scientific) was used to enrich for T cells in single-cell suspension prior to flow cytometry sorting.

Stained cells were analyzed with an LSR Fortessa (BD Biociences) with cell populations defined, gated, and analyzed with FlowJo software (FlowJo, LLC, Ashland, Oregon). Immune populations were identified by forward and side scatter followed by doublet discrimination of viable cells and a sorting strategy modified from [7, 8, 28, 31, 32] depicted in **Supplemental Figure 1A**.

### Ex vivo Treg Isolation and Intracellular Staining

Single cell suspensions were prepared from lungs harvested from mice at the indicated time points as described above. Red blood cells were removed with ACK lysis buffer per the manufacturer’s instructions. Tregs were FACS purified based on the gating strategy depicted in **Supplemental Figure 1C** and plated on 24-well cell culture plates precoated for 2 hours at 37°C with 5 ug/ml of αCD3 (ThermoFisher) and 5 ug/ml of αCD28 (ThermoFisher) at a concentration of 2-5 x 10^5^ cells/ml overnight at 37°C. Adherent cells were then treated with Cell Stimulation Cocktail plus protein transport inhibitor (eBioscience) for 4 hours at 37°C. After incubation, cells were resuspended in PBS and stained with viability dye and fluorochrome conjugated antibodies. Intracellular Areg staining was performed using the Foxp3 transcription factor staining buffer set (eBiosciences) while staining simultaneously for transcription factors – Gata3, Tbet and Rorgt. Areg was the detected by staining with biotinylated goat polyclonal anti-mouse AREG antibody and APC-conjugated streptavidin (1:200) (Biolegend, San Diego, CA).

### Lung Histology and Immunofluorescence Staining

Whole lungs were fixed by tracheal instillation of 10% neutral buffered formalin (MilliporeSigma) at a constant pressure of 25 cm H_2_O as described [33, 34]. Embedded 6 μM sections were cut and stained with Hematoxylin & Eosin (H&E) by the Comparative Pathology Core Laboratory at the School of Veterinary Medicine at the University of Pennsylvania. Slides were scanned using an Aperio ScanScope Model: CS2 (Leica, Wetszler, Germany) at 40X magnification and representative areas were captured and exported as TIF files and processed in Adobe Illustrator.

Immunofluorescence staining of lung sections was performed as described [35] using combinations of primary antibodies in **Supplemental Table 3** including an in-house rabbit anti-proSP-C [36] and other antibodies visualized with fluorescent or HRP conjugated secondary antibodies (**Supplemental Table 3**). Nuclei were identified using DAPI or Hoechst staining. Imaging was performed on an Eclipse 80i upright microscope (Nikon Melville, NY) and processed using ImageJ (Bethesda, MD).

### Picrosirius Red (PSR) Staining

Lung sections were stained for fibrillar collagen using the Picrosirius Red Kit following the manufacturer’s instructions (Polysciences, Inc., Warrington PA). Digital morphometric measurements were performed on captured images from multiple lobes at multiple levels with ten random peripheral lung images devoid of large airways per slide analyzed at a final magnification of 20× using Image J as published [26, 37, 38]. The mean area of PSR staining in each lung field from each section was calculated and expressed as a percentage of total section area as adapted from Henderson, et al [39].

### Murine Pulmonary Function Testing

Lung physiological parameters were assessed in experimental mice anesthetized with intraperitoneal pentobarbital using a Flexivent (SCIREQ, Inc. Toronto Canada) as we described previously [26, 37, 40]. Static lung compliance was determined with the manufacturer’s software using a 2 second breath pause maneuver.

### Bronchoalveolar Lavage Fluid (BALF) Collection and Processing

BALF collected from mice using sequential lavages of lungs with five X 1 ml aliquots of sterile saline was processed for analysis as described [30, 34]. Cell pellets obtained by centrifuging BALF samples at 400 × *g* for 6 minutes were re-suspended in 1 ml of PBS, and total cell counts determined using a NucleoCounter (New Brunswick Scientific, Edison, NJ). Total protein content of cell free BALF was determined using the DC Protein Assay Kit (BioRAD, Inc, Hercules CA) with bovine serum albumin as a standard according to the manufacturer’s instructions.

### RNA Isolation and Quantitative Real Time Polymerase Chain Reaction

RNA was extracted from homogenized lung or isolated GFP+Tregs cells using RNeasy Mini and Micro Kit (Qiagen, Valencia, CA) following the manufacturer’s protocol. The concentration and quality of extracted RNA from the lung tissues were measured using NanoDrop® One (Thermo Scientific, Wilmington, DE) and reverse-transcribed into cDNA using either Taqman Reverse Transcription Reagents (Applied Biosystems, Foster City, CA) or Verso cDNA Synthesis Kit (ThermoFisher).

Quantitative real time PCR (qRT-PCR) for whole lung or isolated Foxp3-GFP+ Treg cell populations was performed as described previously[35]. Transcripts for *Col3A*, *Col2*, *Ki67*, *Topoisomerase2a*, *Areg*, *Ccl2, Ccl17* were amplified using TaqMan Gene Expression Assays in an Applied Biosystems ViiA 7 real-time PCR system with a 384 well plate using RNA primer sequences for all mouse genes listed in **Supplemental Table 2**. Results were normalized to *18S* or *Actb* as indicated.

### RNA Sequencing Processing and Gene Ontogeny Analysis

Analysis of our previously published mouse data set (GSE234604)[29] as well as a published human data set (GSE227136) [41] were performed to interrogate regulatory T-cell biology in murine and human fibrosis. Previously processed data was reclustered using the Leiden algorithm with scanpy.tl.leiden [42].

We identified cell populations using known canonical marker genes or by assessing cluster-defining genes based on differential expressions. Human data set GSE227136 was previously annotated with disease status and our analysis was limited to what these authors previously defined as control and pulmonary fibrosis samples. Finally, receptor ligand interactions were identified using CellChat [43, 44].

### Mouse Organoid Cultures

Organoid culture assays were performed as previously described [45]. AT2 cells were flow sorted as described above, GFP positive fibroblasts were collected via flow cytometry from B6.129S4-Pdgfratm11(EGFP)Sor/J mice (**Supplemental Figure 1A**). In each technical replicate, 5000 AT2 cells were combined with 50,000 *Sca1*^+^/Sca1^-^ *Pdgfra*^+^ lung fibroblasts in 50% Matrigel (Corning; Corning, NY) and 50% Small Airway Growth Medium (SAGM) (Lonza, Basel, Switzerland) in a Falcon Cell Culture Insert. Cell/matrigel suspension solidified and SAGM medium was then added into the bottom of the well. SAGM was prepared according the BulletKit (Lonza, Basel, Switzerland) per the manufacturer’s instructions with some modification (i.e. Hydrocortisone, BSA, Triiodothyronine, EGF, and Epinephrine were omitted). Medium was changed every other day. 10 μM rock inhibitor (Y-27632, MilliporeSigma) was added to the medium for the first two days of culture. Organoids were imaged on an EVOS FL Microscope and were quantified via ImageJ (NIH Bethesda, MD) using “analyze particles” macro.

Organoid cultures for immunohistochemistry were fixed in 4% PFA for 10 minutes then transferred into 50 µl fresh 100% Matrigel (Corning) and the admix mounted (10-20 µl) directly onto glass slides and incubated at 37°C for 30 minutes to allow for polymerization. Samples are then stained and imaged as above.

### Statistics

All data are depicted with dot-plots and presented as group mean ± SEM unless otherwise indicated. Statistical analyses were performed with GraphPad Prism (San Diego, CA). Student’s t-test (1 or 2 tailed as appropriate) were used for 2 groups; Multiple comparisons were done by analysis of variance (ANOVA) was performed with post hoc testing as indicated; survival analyses were performed using Kaplan Meier with Mantel-Cox correction. In all cases statistical significance was considered at p values ≤ 0.05.

## RESULTS

### Ontogeny of Regulatory T Cells During Lung Fibrogenesis

The dynamics of Treg lung infiltration in fibrogenesis were assessed using a murine model of intrinsic PF driven by expression of a clinically relevant mutation in the surfactant protein-C gene (*SFTPC*) identified in spontaneous and familial IPF patients [23–25]. Induction of mutant SP-C expression in adult I^ER^-*Sftpc^I73T^* mice using two sequential doses of oral Tamoxifen (Tam) (**Figure 1A**) produced an elevation in total BALF protein within 1 week of induction (**Supplemental Figure 1A**) accompanied by a dynamic alveolitis with total BALF cells peaking at 2-3 weeks (**Supplemental Figure 1B**) reflective of prior studies [26, 29]. Using flow cytometry and a gating strategy for lung cell suspensions shown in **Supplemental Figure 2**, we identified marked changes in immune cell populations in the SP-C^I73T^ lung after Tam induction that included an increase in tissue neutrophils at 2 weeks and eosinophils at 4 weeks post-injury (**Supplemental Figure 1C-D**).

**Figure 1:**
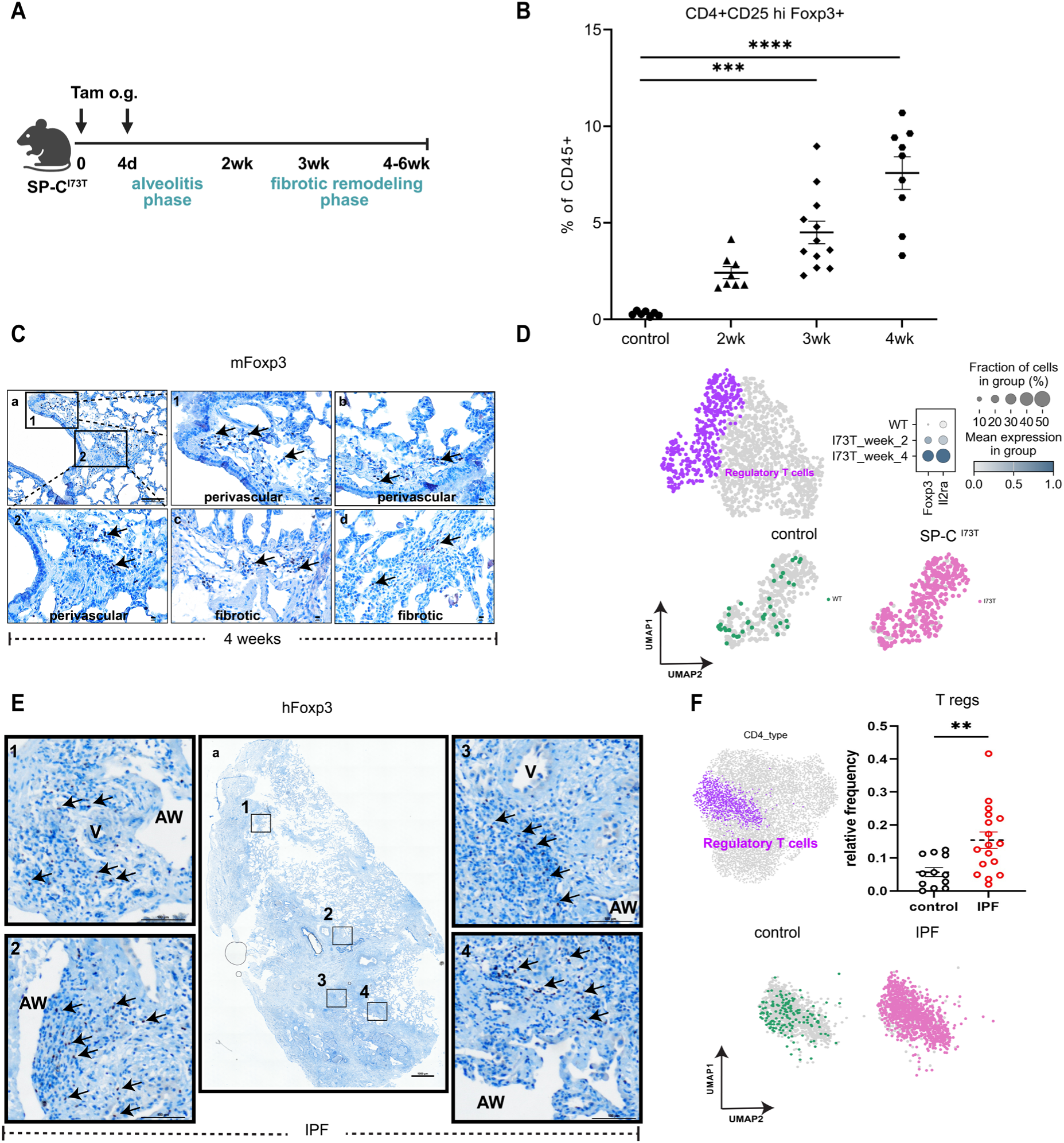
Ontogeny of regulatory T cells in lungs during fibrosis. **(A)** The previously described SP-C^I73T^ mouse pulmonary fibrosis model [26] was utilized with an induction protocol containing 2 sequential doses of tamoxifen delivered by oral gavage [29] that produces 2 distinct phases of overt disease development - alveolitis or peak inflammation (1-2 weeks) and fibrogenesis or fibrotic remodeling (2-6 weeks); **(B)** Lungs from control and SP-C^I73T^ mice were analyzed for Foxp3+Tregs by flow cytometry at indicated time points post Tam induction. Data pooled from three independent experiments with a total of 3 - 12 mice in each group as presented in the scatter plots. Kruskal-Wallis multiple comparison one-way ANOVA was performed. *** p=0.0006 and **** p<0.0001; **(C)** Representative immunohistochemical staining of paraffin-fixed SP-C^I73T^ lungs obtained 4-weeks post induction and stained for murine Foxp3 (*mFoxp3*) expression. Arrows indicate mFoxp3+ Tregs localized in perivascular and fibrotic areas, high power magnification of insets 1,2. Bar: a =100 μm, b-d=10 μm,1-2=10 μm; Images are representative of 3-5 animals per group; **(D)** Uniform manifold approximation and projection (UMAP) analysis of 1,341 CD3+ T cells from lung scRNAseq data (GSE234604) reveals a Foxp3+ Treg cluster following mutant SP-C expression. Gradient dot plots show expression of Treg-specific genes *Foxp3* and *Il2ra* in SP-C^I73T^ mice at 2- and 4-weeks post tamoxifen induction (n=2 per genotype per time point); **(E)** Representative immunohistochemical staining for human Foxp3 (*hFoxp3*) in a paraffin embedded section from a human IPF lung explant. Insets 1-4 are magnified views of boxed regions. Bar 1-4 =100 μm; a=1000 μm **(F)** UMAP visualization of 8,404 CD3+CD4+ T cell subtypes from lung cells analyzed by scRNAseq in accession number GSE227136, comparison of relative frequency of Treg (Foxp3+) clusters between control (n=12) and IPF (n=17) patients, a threshold cut-off > 50 cells was used for analysis; ** p=0.004 by two-tailed Mann-Whitney test.

Inflammation resolution and a transition to fibrogenesis was accompanied by a temporal increase in CD4+CD25^hi^FoxP3+ Tregs reaching 8–10% of all lung CD4 cells by 4 weeks post-Tam treatment (**Figure 1B**). Immunohistochemical staining for Foxp3+ in murine SP-C^I73T^ lung sections 4 weeks post-injury demonstrated Tregs to be localized in both perivascular and fibrotic regions (**Figure 1C**).

To corroborate flowcytometric and IHC analyses, we performed single cell RNAseq (scRNAseq) analysis of cell suspensions from lungs harvested from induced I^ER^-*Sftpc^I73T^* and control (C57/Bl6 “wild-type”) mice. Using marker genes Foxp3 and Il2ra to annotate Tregs, we profiled 1,341 T cells from I^ER^- *Sftpc^I73T^* mice at 2- and 4-weeks post-induction as well as (n= 2 mice per time point) cells from control mice (n=2). A time-dependent increase in Treg-identifying genes was observed in I^ER^-*Sftpc^I73T^* lungs (**Figure 1D**). To translate these findings, we next stained human IPF lung sections for Foxp3. Similar to I^ER^-*Sftpc^I73T^*mice, we found Foxp3+ Tregs adjacent to the vasculature in damaged areas of the lung (**Figure 1E**). Finally, when published human scRNA-seq data from accession number GSE227136 [41] was reanalyzed to compare subsets of CD4+ T cell clusters (**Supplemental Figure 3**), an increase in Treg frequency was detected in samples from IPF patients was observed (**Figure 1F**).

#### Regulatory T cells in Fibrotic Lungs Have Increased Expression of Effector and Activation Markers

Phenotypic characterization of regulatory T cells in SP-C^I73T^ expressing mouse lungs during disease evolution was further characterized using a SP-C^I73T^-Foxp3^eGFP^ line. We first defined the ontogeny of aberrant injury-repair responses in this second line subjected to the same split Tam dosing protocol (**Figure 2A**) and found that SP-C^I73T^-Foxp3^eGFP^ phenocopied the published founder line [29] with similar degrees of weight loss (10-25% of initial body weight) and elevations in BALF cell count and total protein (**Supplemental Figure 4A-C**). We also observed similar kinetics of Treg infiltration post-induction using flow cytometry wherein CD4+Foxp3-GFP+ Tregs were found to peak at 6-8% by 3-4 weeks post Tam induction (**Figure 2A-B**). CD4+Foxp3-GFP+ Tregs sorted from T cell-enriched lung cell suspensions demonstrated increased expression of cell surface molecules associated with inhibitory and effector functions. By 4 weeks post-induction CD4+Foxp3-GFP+ Tregs had elevated expression of CD44, PD-1, CD103, Glucocorticoid-induced tumor necrosis factor-related protein (GITR), T-cell immunoreceptor with immunoglobulin and ITIM domains (TIGIT) compared to Tregs from control, non-uninduced SP-C^I73T^-Foxp3^eGFP^ animals (**Figure 2C-F**).

**Figure 2:**
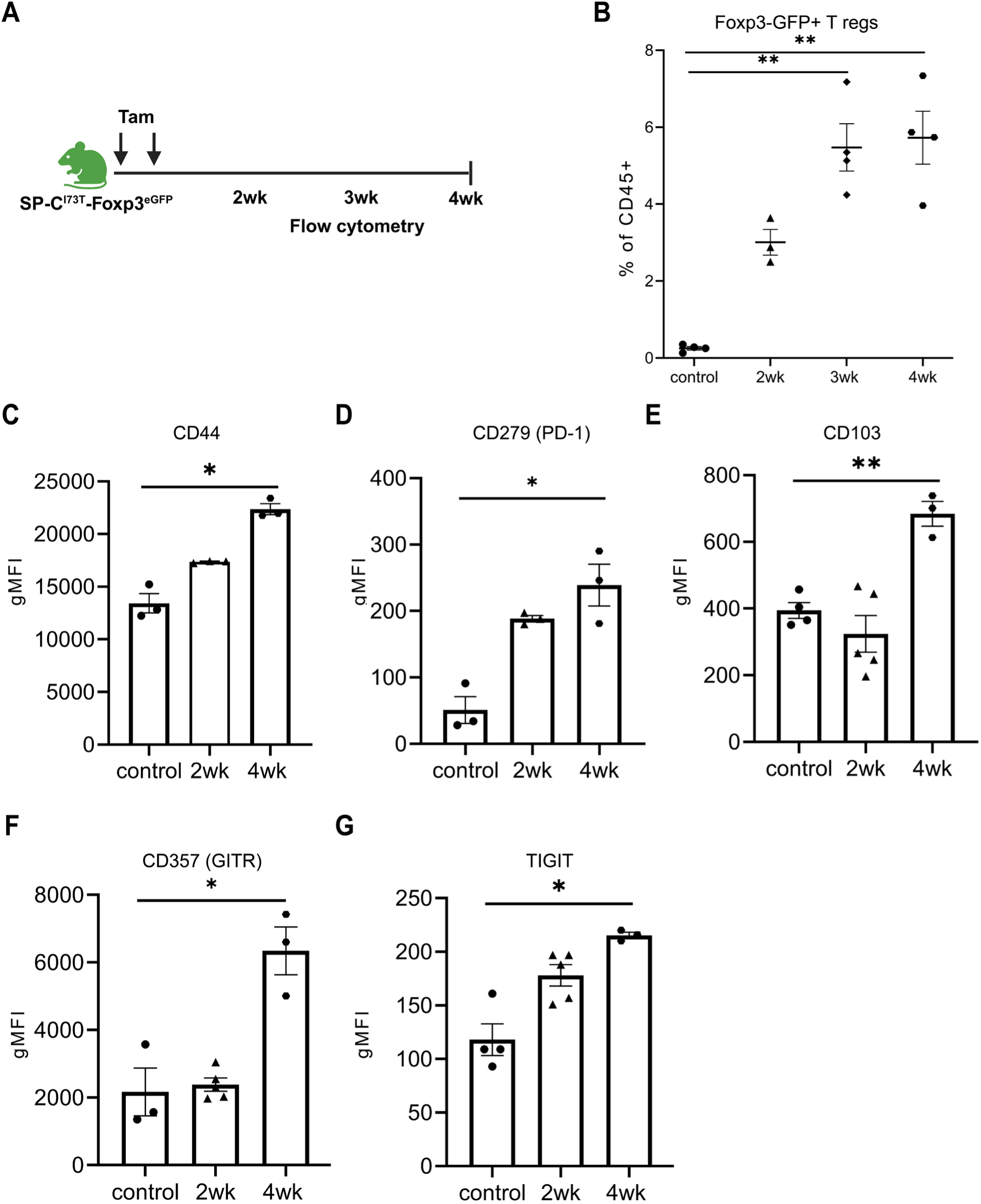
Regulatory T cells in fibrotic lungs have increased expression of effector and activation markers. **(A)** Foxp3^eGFP^ reporter mice were crossed with previously published SP-C^I73T^ mice to generate SP-C^I73T^- Foxp3^eGFP^ mice; **(B)** CD4+Foxp3-GFP+ T regs were analyzed by flow cytometry in lungs harvested at 2, 3 and 4 weeks post Tamoxifen induction of mutant SP-C. Ordinary one-way ANOVA was performed. ** p< 0.005 (n=3-5 mice in each group); **(C-G)** Control and SP-C^I73T^- Foxp3^eGFP^ lungs were digested, and samples were enriched for T cells using magnetic bead depletion as described in *Materials and Methods*. Flow cytometric analysis and quantification of geometric mean fluorescence intensity of effector and activation markers expressed by CD4+Foxp3-GFP+ T regs in lung tissue of SP-C^I73T^- Foxp3^eGFP^ mice 2 and 4-weeks post mutant *Sftpc* induction: **(C)** CD44; **(D)** CD279 (PD-1); **(E)** CD103; (**F)** GITR; **(G)** TIGIT. For each marker, ordinary one-way ANOVA was performed and *p<0.05 and ** p< 0.005, n = 3 - 5 mice per group.

### Treg Depletion After Injury Promotes Enhanced Fibrogenesis in I^ER^-Sftpc^I73T^ Mice

Having identified a peak in activated lung Tregs coinciding with transition to a fibroproliferative phase in the model [26, 29] (**Figure 1A**), we next asked whether Treg depletion initiated at this epoch (2-3 weeks post-induction) would alter fibrotic remodeling. For this we adopted a genetic approach and generated a SP-C^I73T^-Foxp3^DTR^ mouse line to allow for temporally controlled, systemic Treg ablation using diphtheria toxin (DT) administration. We first assessed DT dosages used in prior published studies to interrogate LPS and flu induced models of lung injury [18, 46] but found that a dosing schedule of 50, 10, and 10 µg/kg beginning at Day 12 post-Tam induction of the SP-C^I73T^-Foxp3^DTR^ line led to unacceptable weight loss (30%) (**Supplemental Fig. 5A**). Based on efficacy data generated in uninduced SP-C^I73T^-Foxp3^DTR^ mice showing effective Treg depletion in the spleen within 48 hours of a single dose of DT (**Supplemental Fig. 5B**), we adopted a modified dosing protocol of 5 µg/kg delivered every 48 hours which avoided increased morbidity (additional weight loss beyond SP-C^I73T^ induction (**Supplemental Fig. 5C**) or increases in BAL cell counts (**Supplemental Figure 5D**). while also achieving ∼80% splenic Treg depletion in Tam-induced SP-C^I73T^- Foxp3^DTR^ mice at 1- and 2-weeks post-induction. After optimizing dosing, we administered this modified DT dose starting at 17 days post-Tam induction and continued through endpoint analysis at day 26 **(Figure 3A).**

**Figure 3:**
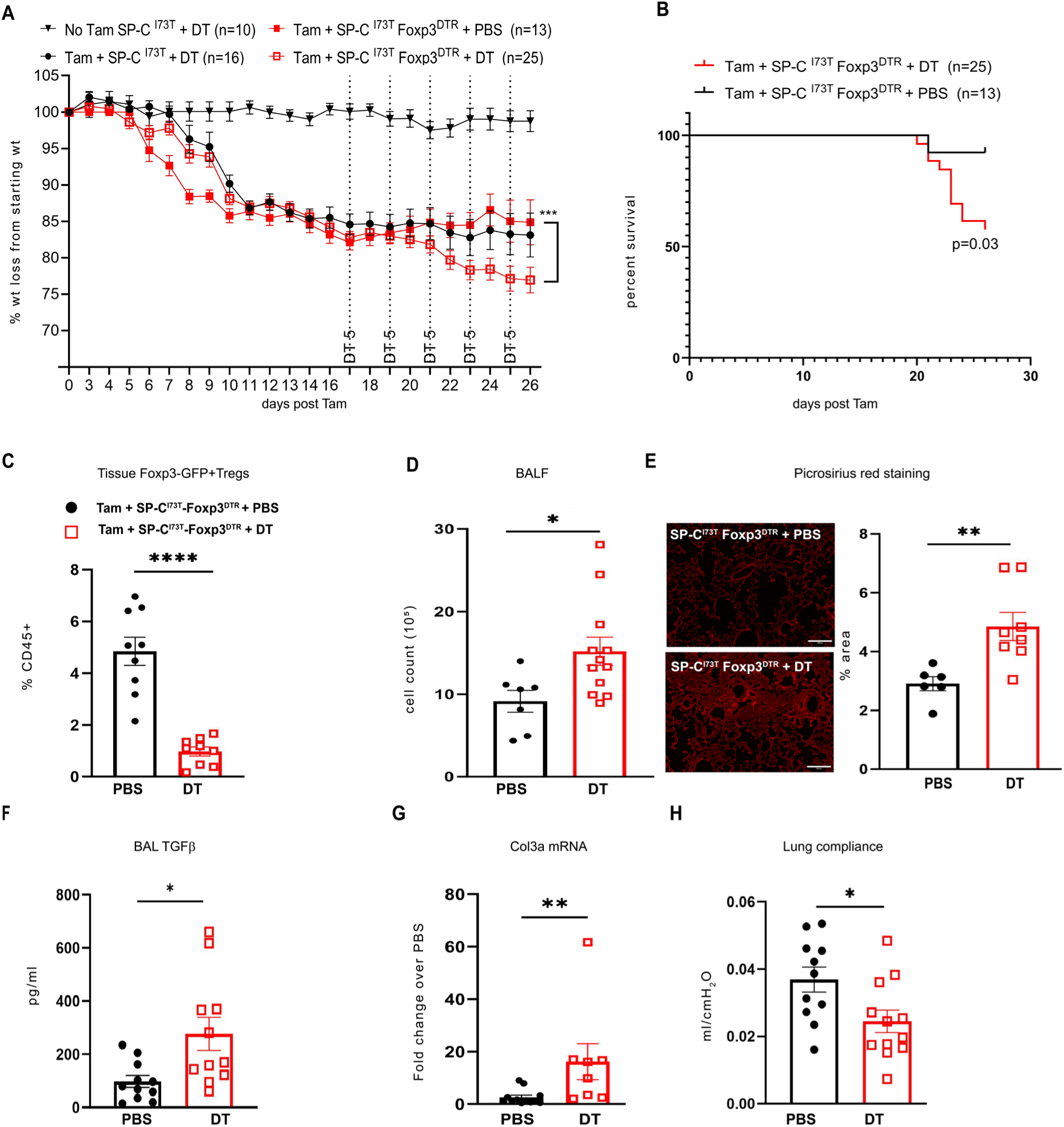
Depleting T regs during fibrogenesis worsens the disease phenotype in I^ER^-SP-C^I73T^Foxp3^DTR^ mice. (**A**) I^ER^-SP-C^I73T^-Foxp3^DTR^ knock-in mice generated as described in *Materials and* Methods nSP-C^I73T^-Foxp3^DTR^ mouse line and AT2 specific lung injury were first induced with Tamoxifen administered via oral gavage on Days 0 and 4. 5μg of DT or PBS was administered i.p. every 48 hours starting at day 17 post Tam induction and weight loss was recorded every day in SP-C^I73T^ and SP-C^I73T^-Foxp3^DTR^ mice. Data are pooled from three independent experiments. ***p<0.0001 by two-way ANOVA (SP-C^I73T^ uninduced - no tam + DT n=10, SP-C^I73T^ + DT n= 16, SP-C^I73T^-Foxp3^DTR^ + PBS n=13, SP- C^I73T^-Foxp3^DTR^ + DT n=27); (**B**) Kaplan Meier survival curves of SP-C^I73T^-Foxp3^DTR^ mice that either received PBS (n=13) or DT (n=25). Log-rank (Mantel-Cox) test was performed with p value 0.0391; (**C**) Lungs from SP-C^I73T^-Foxp3^DTR^ mice receiving DT (n=9) or PBS (n=9) were harvested and cell suspensions analyzed by flow cytometry for GFP+ Tregs. *** p value < 0.0001 by unpaired t-test with Welch’s correction; (**D**) Quantification of total BALF cell counts between SP-C^I73T^-Foxp3^DTR^ mice treated with PBS (n=7) or DT (n=12) 26 days after Tam induction. * p=0.01 by unpaired t-test with Welch’s correction; (**E**)(*Left*) Representative Picrosirius red stained fields from lung sections from DT or PBS treated SP-C^I73T^-Foxp3^DTR^ mice 26 days after Tam induction. Bar = 200μm. (*Right*) Quantification was performed using ImageJ and data expressed as a percentage area staining for PSR. ** p = 0.004 by unpaired t-test with Welch’s correction; (**F**) BALF TGFβ quantified using ELISA (PBS n=11, DT n= 11); (**G**) qRT- PCR analysis of whole lung mRNA for *Col3a* (PBS n=12, DT n= 8); For E- G, scatter plots with individual values data was analyzed by unpaired t-test *p<0.05 ** p<0.01; (H) Box and Whiskers plot of static lung compliance (Cst) measured using Flex-Vent as described in *Materials and Methods* (PBS n=11, DT n= 12). Individual values for each animal are depicted and analyzed by unpaired t-test *p<0.05 ** p<0.01 vs PBS.

Consistent with a protective and / or reparative role of Tregs in other lung injury models [17, 19, 20], Treg depletion with Diptheria toxin worsened morbidity and mortality in SP-C^I73T^ - Foxp3^DTR^ mice causing increased weight loss (**Figure 3A)** and decreased survival (**Figure 3B**) compared to controls (Tam induced SP-C^I73T^- Foxp3^DTR^ mice receiving PBS). Since both control groups (either Tam induced SP-C^I73T^- Foxp3^DTR^ mice receiving PBS or Tam induced SP-C^I73T^ receiving DT) showed similar BALF cell counts at day 26, suggesting comparable injury and inflammation, SP-C^I73T^-Foxp3^DTR^ mice receiving PBS served as controls for subsequent experiments.

Flow cytometric analysis confirmed significant Treg depletion in the lungs of DT-treated mice at 26 days (**Figure 3C**) with Treg-depleted SP-C^I73T^ - Foxp3^DTR^ mice developing more severe alveolitis (**Figure 3D**). Commensurate with this finding, there was also increased expression of Ccl2 and Ccl17, cytokines involved in monocyte and eosinophil trafficking to the lungs in the SP-C^I73T^ model (**Supplemental Figure 6**). Most significantly, Treg-depleted SP-C^I73T^ - Foxp3^DTR^ mice developed increases in key fibrotic endpoints. Histological analysis of lung section stained with Picrosirius red stain showed increases in fibrillar collagen throughout the lung regions (**Figure 3E**) with higher levels of the pro-fibrotic cytokine TGF- β **(Figure 3F**). Simultaneously we found higher expression of *Col3a* (**Figure 3G**) coincident with acquisition of restrictive lung physiology as demonstrated by a decrease in static lung compliance (**Figure 3H**). In contrast, a preventative intervention strategy in which Diphtheria toxin mediated deletion of Tregs was performed prior to Tamoxifen induction of mutant *Sftpc* failed to alter the pathogenic course (**Supplemental Figure 7)**. In summary, Treg depletion initiated at the transition to fibrogenesis exacerbates pro-fibrotic and pro-inflammatory responses to SP-C^I73T^ expression.

### Tregs in SP-C^I73T^ mice are Type 2 Skewed Post-injury

It is widely accepted that in addition to classification based on surface and activation markers Tregs can be subdivided into subsets based on injury type, inflammatory cytokines, and signals from structural cells at the injury site [13, 47, 48]. We performed additional analyses of our published single-cell RNA sequencing dataset [29] derived from lung samples at 2- and 4-weeks post-Tam induction to assess transcriptional differences in T cell subsets. Single cell suspensions of lungs from Tam treated SP-C^I73T^ and control (uninduced) mice, were profiled for CD45+ cell populations. Using lineage-specific markers and published gene signature, we classified T cells into four clusters: CD4+, CD8+, and two Foxp3+ (Treg) subsets which were further subdivided based on the level of proliferation markers (**Figure 4A**). At both 2- and 4-weeks post-induction, the majority of the CD3(+) T cell clusters were composed of cells contributed by SP-C^I73T^ animals compared to controls (**Supplemental Figure 8).** Tregs from SP-C^I73T^ mice expressed high levels of the Th2 transcription factor Gata3, while Th1 (Tbx21) and Th17 (Rorc) markers were absent, indicating a Type 2 skewing (**Figure 4B**). Genes previously shown to be regulated by GATA3 (Ilrl1, Maf) [48] were also upregulated at week 4 compared to controls (**Figure 4C**).

**Figure 4:**
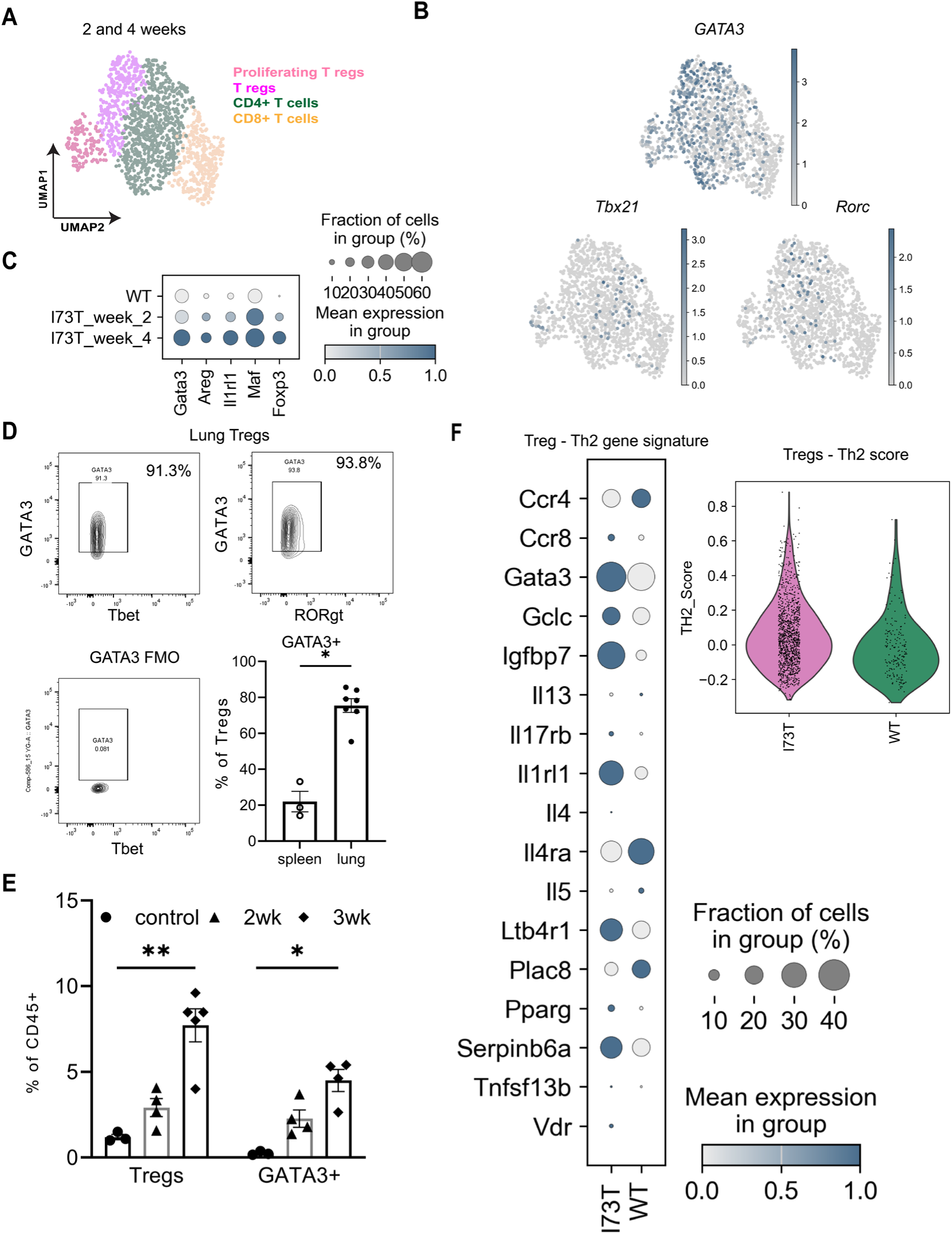
Tregs in SP-C^I73T^ mice are type 2 skewed post injury. **(A)** Reanalysis of a previously published and deposited scRNAseq data set (GSE234604). UMAP clustering of 1,341 T cells in SP-C^I73T^ and uninduced control mouse lungs identified 2 populations of Foxp3+ T regs, CD4+ and CD8+ T cells at 2- and 4-weeks post injury; **(B)** UMAP projection of all T cells identifies *Gata3* as the major transcription factor expressed by Treg populations; **(C)** Gradient plot depicting *Gata3*-regulated genes - in lung Tregs at 2, 4 weeks post-injury and control (uninduced) mice; (**D)** Representative FACS plot of lung cell suspension obtained 3 weeks post-Tam induction showing CD25hi Foxp3+ T regs co-stained for transcription factors - Tbet (Th1), GATA3 (Th2) and Rorgt (Th17) as labeled. Spleens (n=3) from of SP-C^I73T^ animals served as controls. Unpaired t-test was performed and *p=0.01; **(E)** Lungs from control and SP-C^I73T^ mice were analyzed and quantified by flow cytometry for transcription factors – Foxp3 and GATA3 flow cytometry at indicated time points post Tam induction. Ordinary one-way ANOVA was performed. * p< 0.05 ** p< 0.005. n=3-5 mice per group; **(F)** Comparative analysis of deposited scRNAseq data set (GSE234604) for genes associated with Th2 immune response 4- weeks post Tam induction. Shown are data from control and SP-C^I73T^ mice represented in a gradient plot. Genes from the gradient plot were combined to generate a Th2 gene score calculated for T regs from SP-C ^I73T^ and uninduced control mouse lungs.

To corroborate the computational analysis, we then stained lung and spleen single-cell suspensions from SP-C^I73T^ mice for Treg markers and Th1, Th2, and Th17 transcription factors. Flow cytometry confirmed that 70-80% of lung Tregs were GATA3+ (**Figure 4D**). In parallel with the increase in lung Foxp3+ Tregs observed post injury, GATA3+ cells were also significantly elevated at 3 weeks post injury in SP-C^I73T^ compared to uninjured control lungs (**Figure 4E**). Additionally, using a published Th2 signature gene set [49], lung Tregs from Tam-induced SP-C^I73T^ mice showed an increased Th2 score compared to controls (**Figure 4F**). In total, both scRNA-seq and flow cytometry demonstrate that Tregs in the lungs of induced SP-C^I73T^ mice are more Type 2 skewed than those from uninjured controls.

### T regs in SP-C^I73T^ Mice Are Proliferative and Generate Pro-Reparative Queues

We next assessed if lung Tregs from injured SP-C^I73T^ mice could produce factors essential to restore alveolar homeostasis. Lung CD4+GFP+ Tregs were collected using the sorting strategy shown in **Supplemental Figure 1C** from Tam-induced Foxp3^EGFP^ - SP-C^I73T^ mice at weeks - 2, 3, and 4 and total RNA extracted (**Figure 5A**). Consistent with the scRNAseq data (**Figure 4A)**, qRT-PCR analysis of Foxp3- GFP+Tregs showed increased expression of replication markers – *Ki67* and *Topoisomerase2a* at 2 weeks post injury, indicating appearance of a proliferative Treg population early after induction (**Figure 5B-5C**). Later in the course (4 weeks post *Sftpc^I73T^*induction), isolated Foxp3-GFP+Tregs were found to have increased mRNA expression of amphiregulin (*Areg), a* pro-reparative growth factor (**Figure 5D**). Additionally, flow cytometric analysis from this same time point indicated that GATA3 expressing lung Foxp3-GFP+ Tregs also produce Areg protein (**Figure 5E; Supplemental Figure 9A**) which was accompanied by a 5-6 fold increase in amphiregulin content in BAL fluid from injured cohorts (**Figure 5F**). We also identified *Spp1* (Osteopontin) in the Treg transcriptome (**Figure 6C)** as well as increased levels of osteopontin protein in BALF (**Supplemental Figure 9B**).

**Figure 5:**
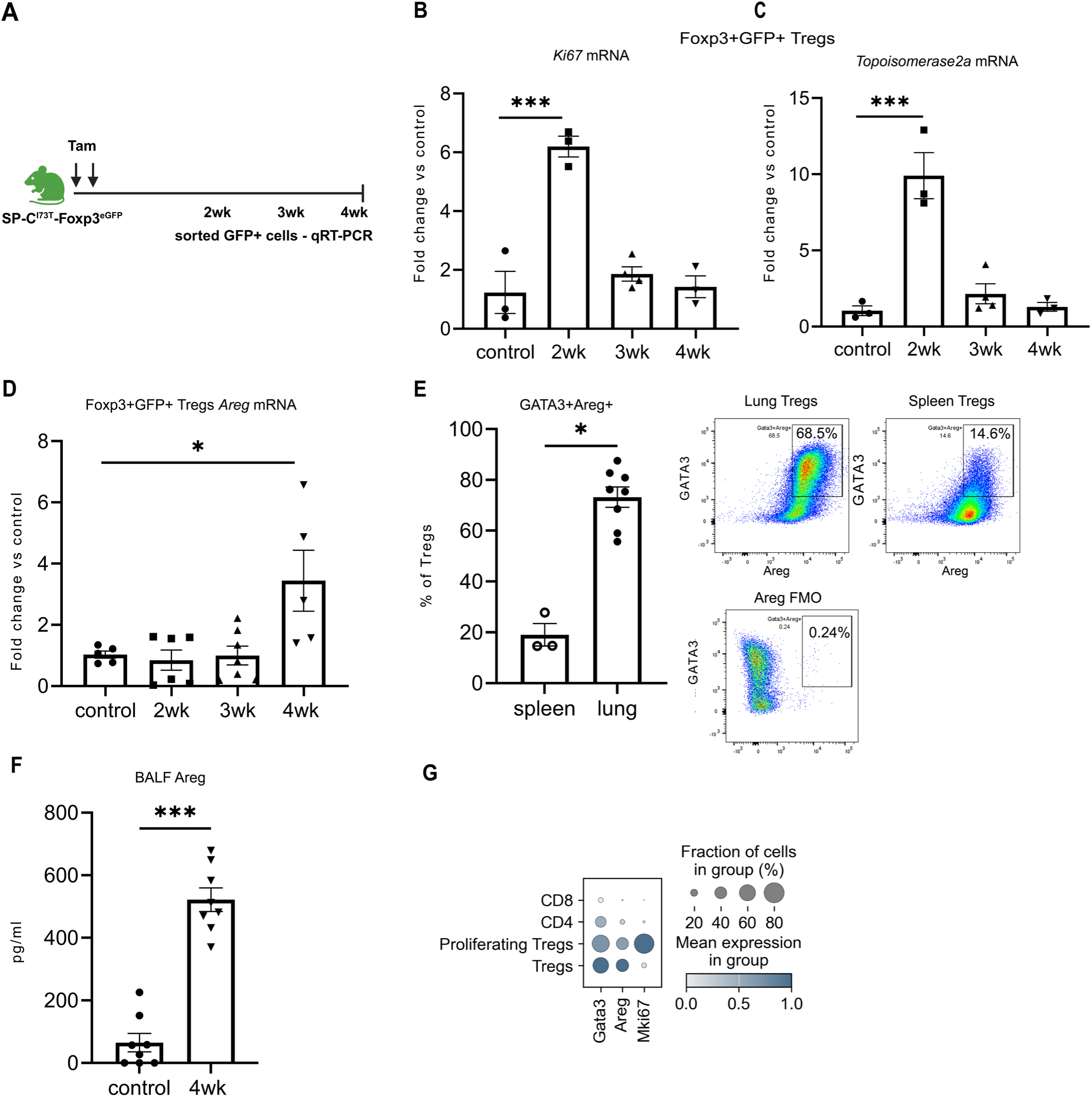
Tregs in SP-C^I73T^ mice are proliferative and make pro-reparative factors. **(A)** Schematic for two dose Tamoxifen induction (days 0 and 4) of I^ER^-SP-C^I73T^-Foxp3^eGFP^ mice. Foxp3-GFP+ Tregs were FACS purified at 2-, 3- and 4-weeks post Tam induction and total RNA isolated. **(B-D)** Scatter plots of qRT-PCR analysis of Treg mRNA for expression of: (**B**) *Ki67*; (**C**) *Top2a(*Topoisomerase2a); (**D**) *Areg*. For each gene ordinary one-way ANOVA was performed. * p< 0.05 *** p< 0.0001. n=3-7 mice per group; **(E)** Foxp3-GFP+ Tregs were FACS sorted from lungs and spleen of SP-C^I73T^mice 4 weeks post Tam induction and plated overnight. Cells were then treated with a cell stimulation cocktail that includes Brefeldin A, a protein transport inhibitor for 4 hours, fixed and stained for GATA3 and intracellular amphiregulin (Areg). Identified Gata3+/Areg+ cells were expressed as a percentage of total Tregs. Comparisons were made using an unpaired t-test. *p=0.01 n=3-8 mice per group; **(F)** Areg levels of BALF collected 4wks after Tam induction and quantitated by ELISA. ***p=0.0002 by unpaired t-test; n= 7-8 per group; **(G)** Gradient plots from the previously published scRNA-seq dataset GSE234604 were re-analyzed to compare the expression of *Areg, GATA3 and Ki67* among Tregs, proliferating Tregs, CD4⁺, and CD8⁺ T cells identified in single cell suspensions prepared from lung tissue 2- and 4-weeks post-Tam induction.

**Figure 6:**
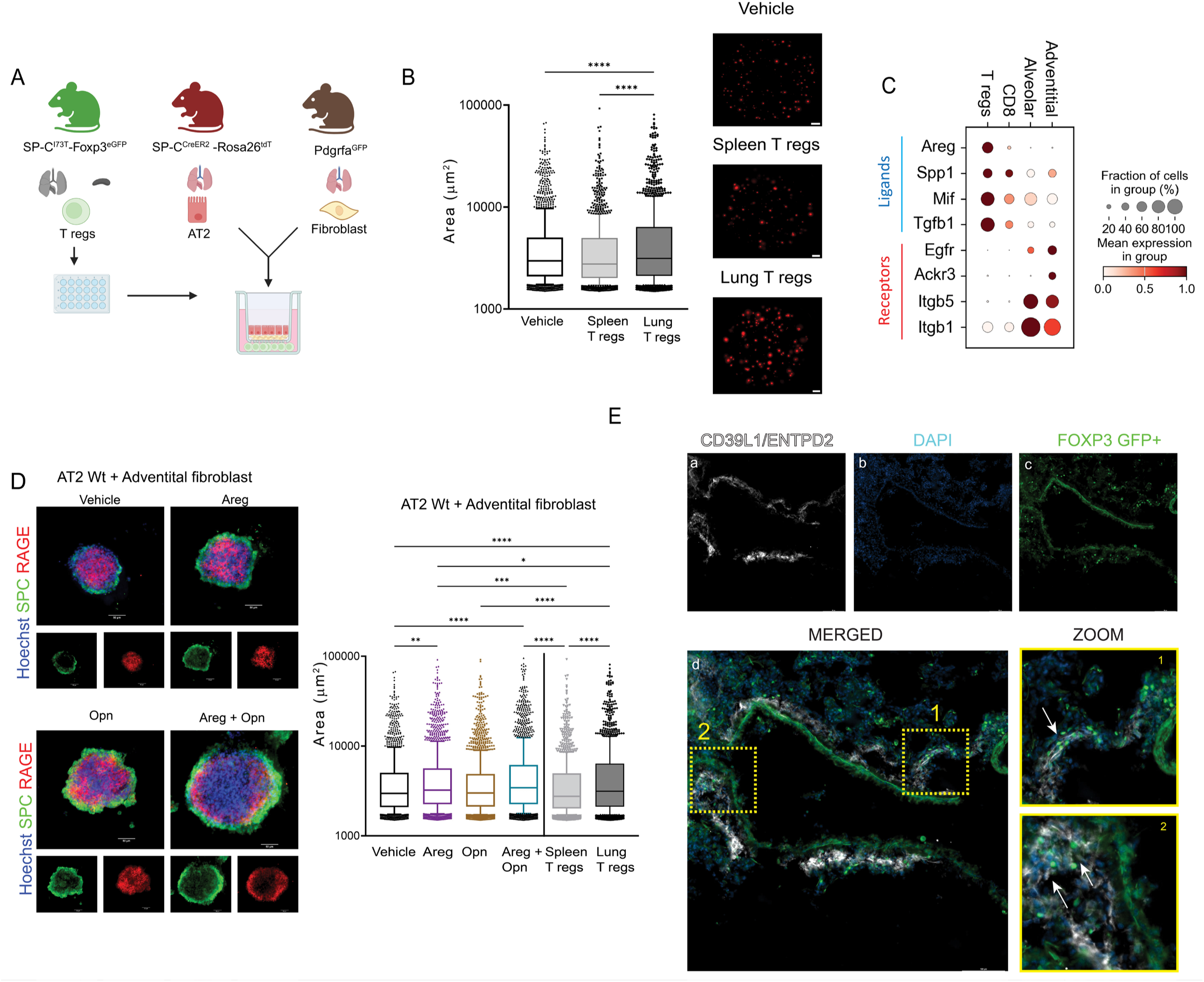
Lung SP-C^I73T^ T regs augment alveolar organoid growth supported by adventitial fibroblasts in vitro. **(A)** Schematic depicting tri- cellular organoid cultures generated using lung Tregs derived from SP-C^I73T^-Foxp3^eGFP^ mice 3-4 weeks after Tam induction, AT2 cells FACS sorted from SP-C^CreER2^-Rosa26^Tdt^ mice, and adventitial fibroblasts isolated from Pdgrfa^GFP^ mice; (**B)** Tricellular organoids were cultured for 2 weeks with activated CD3+CD28 Tregs replenished every week. Organoid cultures were imaged using EVOS FL Auto and images were quantified using an ImageJ an “analyze particle” macro with minimum area threshold set to1500 μm^2^. The experiment was independently repeated at least three times, each with three biological and three technical replicates. ****p<0.00005 done by Ordinary one-way ANOVA; **(C)** Gradient plot from a previously published scRNA-seq dataset (GSE234604) re-analyzed for expression of potential ligands made by immune cells as labeled (CD8+ T cell or Foxp3+ T regs) and cognate receptor expression on two fibroblast sub-populations (adventitial or alveolar) as labeled that are each increased in SP-C^I73T^ mouse lungs at 2- and 4-week post Tam induction; **(D)** Bi-cellular organoids containing FACS purified TdTom+ AT2 and ScaI+ adventitial fibroblast were cultured with either 100 nM Areg, 200 nM Opn, or Areg + Opn for 2 weeks. For comparison, tricellular organoids containing lung or spleen Tregs were prepared and co-cultured for 2 wks. (Left) Representative immunofluorescence staining of fixed wholemount organoid cultures co-stained for proSP-C and RAGE as labeled. Bars 50 μm; (Right) Live organoid cultures were imaged as in (B) (**E)** Representative immunofluorescence staining of fixed, frozen lung sections prepared form SP-C^I73T^-Foxp3 ^eGFP^ mice at 26 days post Tam induction stained for (a) adventitial fibroblasts as CD39L1+ (white); (b) nuclei as DAPI+ (blue); (c) inherent Foxp3+GFP+ fluorescence identifying Tregs (green); (d) composite overlay. Bars 100 μm. Insets 1 and 2 are magnified views of the boxed perivascular regions of lung with GFP+ Tregs and CD39L1+ adventitial fibroblasts.

Finally, returning to our scRNAseq data, we corroborated these findings by showing at the transcriptomic level, compared to conventional CD4+ and CD8+ T cells, both proliferating Foxp3+Tregs from early time points as well as Tregs from late time points were GATA3+ and capable of producing amphiregulin (**Figure 5G**).

#### Lung SP-C^I73T^ T regs Augment Adventitial Fibroblast Supported Alveolar Organoid Growth in vitro

To functionally characterize a role for Tregs in the distal lung niche post-injury, we modeled epithelial-mesenchymal-immune crosstalk by establishing a tri-cell 3D co-culture organoid assay containing isolated AT2 cells, lung mesenchymal populations, and Foxp3-GFP+Tregs (**Figure 6A)**. We focused on receptor-ligand interactions between Tregs and two major fibroblast populations – adventitial and alveolar, previously reported as key drivers of fibrotic remodeling in SP-C^I73T^ model of fibrosis [29]. Based on our prior bicellular 3D co-culture studies, adventitial fibroblasts were markedly superior to alveolar fibroblasts in supporting AT2 derived alveolar organoids [29]. We first employed Foxp3-GFP+Tregs from both lung and spleen of Tam-induced Foxp3^EGFP^ - SP-C^I73T^ mice specific with FACS isolated Td-Tomato (+) AT2 cells, cultured with adventitial fibroblasts for 2 weeks. Interestingly, lung- but not splenic- Tregs promoted larger organoid size, suggesting tissue-specific Tregs enhance lung epithelial-mesenchymal niche repair (**Figure 6B**). We confirmed that only adventitial, not alveolar, fibroblasts supported AT2-derived alveolar organoid formation in a tri-cell 3D co-culture system (**Supplemental Figure 10**).

We next assessed the capacity of individual candidate ligands from Tregs to support similar reparative functions. We first interrogated our scRNAseq data to identify potential interactions between mesenchymal and T cell clusters. **I**n addition to amphiregulin, we also identified Treg specific upregulation of transcripts for two other growth factors, Spp1 (Osteopontin) and Mif in SP-C^I73T^ mice 4 weeks post injury accompanied by increased expression of EGF receptors – Itgb1, Itgb5 and Egfr on adventitial and alveolar fibroblast populations (**Figure 6C**). We then assayed, in the same 3D system, the ability of Areg and Spp1, alone or in combination to enhance organoid formation. As shown in **Figure 6D**, in co-cultures with AT2 cells and adventitial fibroblasts, the combination of Areg and Opn mimicked the effect size of Tregs from injured lungs. As previously reported, alveolar fibroblast did not support organoid formation and did not respond to Areg or Spp1 [21]. Lastly, immunostaining of Tam-induced Foxp3^EGFP^-SP-C^I73T^ mouse lungs 4 weeks after injury (**Figure 6E**) revealed that adventitial fibroblasts resided near the vasculature, in a distribution similar to the Tregs. The combination of spatial localization and reductionist culture systems provides support for a role for Treg-mesenchymal crosstalk mediated by Areg and Osteopontin in lung repair.

## DISCUSSION

Idiopathic pulmonary fibrosis is increasingly recognized as a polycellular disease of the distal lung parenchymal niche. While the role of both epithelial and mesenchymal cell populations in the pathogenic cascade is well established, this study has uncovered a specialized role for Tregs during lung fibrogenesis using a spontaneous preclinical mouse model of IPF. The model permitted a precise determination of the ontogeny of Tregs expansion in which we observed a time-dependent accumulation of Tregs peaking during the transition from inflammation at 3-4 weeks post-injury (**Figures 1 and 2**). Tregs in fibrotic lungs exhibited increased expression of activation and effector markers and were skewed toward a GATA-3+, Type 2 phenotype, suggesting a tissue-specific, immunoregulatory program. These Tregs were also highly proliferative early (2 weeks) post injury and expressed pro-repair growth factors including Areg at later points, suggesting a potential role in promoting tissue repair/regeneration which was confirmed by functional assays in which inclusion of Tregs enhanced adventitial fibroblast–supported alveolar organoid growth *in vitro*. Moreover, depletion of Tregs *in vivo* during the resolution phase exacerbated inflammatory and fibrotic outcomes further underscoring their critical contribution to limiting disease severity. Collectively, our data identifies immune–epithelial–mesenchymal crosstalk as a key component of the reparative niche in fibrotic lung tissue. This tri-lineage interaction may be central to understanding the cellular coordination required for fibrosis resolution and tissue regeneration, offering novel mechanistic insight and therapeutic opportunities in IPF.

Regulatory T cells, as a specialized subset of CD4+ T cells, are best known for their role in preventing autoimmunity and maintaining immune homeostasis [9]. Emerging evidence suggests they may also function in tissue repair [20, 21, 50]; however, prior studies in preclinical models of lung fibrogenesis have been less conclusive [31, 51, 52]. Expansion of Treg populations via systemic IL-2 delivery or adoptive transfer of splenic Tregs each performed <u>before</u> bleomycin challenge, worsened fibrogenic endpoints, as evidenced by higher Ashcroft scores, increased trichrome (collagen staining, and reduced survival [31]. In contrast, anti-CD25-mediated Treg depletion initiated a day before bleomycin instillation improved a variety of fibrosis markers [52]. Another study suggested a dual role for Tregs in bleomycin-dependent fibrosis, showing that early depletion (3 days before bleomycin) reduced fibrosis, while later depletion (9–16 days after bleomycin) increased fibrosis [51]. This lack of equipoise has been, in part, due to limitations of current rodent models of fibrosis which rely on exogenous lung injury, specifically delivery of bleomycin, LPS, or radiation. In the current study the use of the mutant Sftpc murine model offered a unique opportunity to both precisely define and better contextualize Treg influences on fibrogenesis. In this model, Tregs were found to appear in significant numbers at the transition from inflammation to fibrogenesis while their depletion at this time point worsened fibrotic endpoints (**Figure 3**). The protective or reparative effect of Tregs was similar to that observed after severe influenza lung infection [21] as well as LPS induced lung injury [17] and has also been observed in a murine model of skin fibrosis [48].

While many prior studies focused on the interactions between Tregs and epithelial cells [17, 18, 51], modulation of fibroblast state is also increasingly recognized as a key event in tissue remodeling after injury with excessive deposition of extracellular matrix (ECM) components by collagen producing fibroblast populations. Extensive application of scRNAseq and bulk gene expression analysis across a variety of preclinical lung models and human samples under both normal (homeostatic) and disease states has now defined a variety of fibroblast populations in the distal lung [29, 53–56]. Adventitial (Pdgfr+; Col14a+, Pi16+) fibroblasts, located near infiltrating Tregs during injury (see [21] and **Figure 6E**) have been shown to enhance colony forming efficiency of AT2 cell derived lung organoids ex vivo [21, 57]. The current work extends this concept wherein the inclusion of Tregs in tricellular 3D organoids containing isolated adventitial fibroblasts and AT2 cells enhanced organoid growth (**Figure 6**). Furthermore, the proliferative effect of Tregs on AT2 cells was lost when alveolar fibroblasts were substituted in this culture supporting recent findings of an adventitial fibroblast dependent circuit [21]. As with previous reports, we see that Areg is a potent driver of this circuit, however this effect is further potentiated by the addition of osteopontin to the organoid culture. This novel observation indicates Tregs produce a complex milieu to stimulate the mesenchyme after injury.

The impact of Tregs on fibrotic remodeling was also niche specific as the enhanced organoid formation was not observed by substitution of spleen derived Tregs adding to recent studies defining the lung specific phenotype of Tregs during injury [48, 57–60]. In our model Tregs are marked by the expression of *Gata3*, a transcription factor established to play a cardinal role in Treg physiology during inflammation [61] but not well defined in the context of lung fibrosis. Our findings suggest that the Gata3 induced Th2 phenotype promotes a pro-reparative state that can, in part, be mechanistically linked to Gata3 activation of Areg expression. Though reports of Treg derived osteopontin have been previously described [15, 62, 63] there is no direct evidence in lung Tregs linking Gata3 to *Spp1* expression. Therefore, our identification osteopontin as a modulator of proliferation in our triculture system suggests that Th2 skewing is not the only driver of the lung Treg niche.

There are several limitations to our study. While the reductionist organoid data is compelling, we could not directly address *in vivo* whether Treg-specific expression of amphiregulin, osteopontin or other growth factors such as KGF, is essential for lung repair and regeneration following AT2 cell-specific injury. To rigorously investigate this, future studies will require the use of conditional knockout models such as Areg fl/fl-Foxp3 Cre mice or adoptive transfer approaches employing wild-type or growth factor-deficient (e.g., *Spp1* or *FGF7* null) Tregs, as previously described in influenza [20, 21] and LPS-induced and left-lung pneumonectomy lung injury models [19]. Unlike prior studies, our work suggests that amphiregulin is not the sole factor essential for epithelial repair, and Tregs are capable of producing multiple growth factors that support adventitial fibroblast-mediated AT2 cell growth *in vitro*. Another important limitation and future direction of this study is to delineate the mechanisms driving Treg trafficking and proliferation in the lung following injury. Epithelial injury may induce elevated expression of IL-33, an alarmin known to promote the expansion of IL-13–producing ST2+Tregs. This subset has been shown to suppress pro-inflammatory cytokines such as IL-6, G-CSF, and MCP-1, thereby limiting the accumulation of Ly6c hi monocytes, as demonstrated in a bleomycin-induced acute lung injury model [64]. Supporting this possibility, Tregs from SP-C ^I73T^ mice exhibit increased expression of *Il1rl1* **(Figure 4F),** implicating a potential role for IL-33/ST2 signaling in the observed Treg response.

Finally, while this study has been focused on the role of Tregs in fibrotic remodeling we recognize that other populations of T lymphocytes could play a reparative role in our model system. Dahlgren et al. [65] have highlighted the importance of group 2 innate lymphoid cells (ILC2s) as key immune regulators within the alveolar niche, particularly in the context of tissue repair and remodeling during type 2 helminth infection. Their work demonstrated that ILC2s localize near Tregs within the lung bronchi and large vessels and are supported by IL-33 and TSLP-producing adventitial fibroblasts that promote their accumulation and activation. While their focus was on ILC2 localization and expansion in response to IL-33 signaling during type 2 immunity, our model of fibrosis reveals a parallel spatial relationship: Foxp3-GFP+ Tregs are found in close proximity to CD39L1/ETPD2+ adventitial fibroblasts (**Figure 6E).** Furthermore, our *in vitro* organoid experiments **(Figures 6B–D)** show that only the combination of adventitial fibroblasts and lung-derived Tregs supports AT2 organoid growth and differentiation. These observations suggest a potentially broader role for adventitial fibroblasts in orchestrating immune-stromal interactions beyond ILC2s, possibly extending to Treg-mediated tissue repair-a hypothesis that warrants further investigation using targeted *in vivo* approaches.

In conclusion, our study underscores the role of the regulatory arm of the immune system in the control of fibroblast activation and regulation of the fibrotic lung niche. Our results suggest that treatment modalities focused on augmenting the function of lung Tregs or functionally mimicking their pro-reparative repertoire may have beneficial effects in treating fibrosis in this organ. In addition, in contrast to targeting a single antifibrotic or profibrotic pathway, we speculate that augmenting cell types that naturally suppress fibrosis or promote niche dynamics skewed towards repair will capitalize on the multiple pronged signals used by these cells to mediate their antifibrotic effects.

## AUTHOR CONTRIBUTIONS

AM and MFB developed the concept. AM, LR, and MFB designed the experiments. YT, PC, SI, KC, CHC and TD performed *in vivo* animal experiments. AM, SB, and LR conducted experiments and analyzed data. WRB and LR performed bioinformatic analysis. JK, AM, and LRR conceived, validated, and optimized flow cytometry strategies. LRR, JK, AM, and MFB interpreted data and generated figures. AM and MFB drafted the original manuscript. LRR, SB, JBK, WRB and MFB edited the manuscript. All authors reviewed and approved the final version prior to submission.

## ACKNOWLEDGMENTS

The authors wish to thank members of the Beers Lab and Anne Sperling, Ph.D. for insightful discussions. MFB is the Robert L. Mayock and David A. Cooper Professor of Medicine. The authors thank the Penn Genomic and Sequencing Core (RRID:SCR_022383). Flow cytometry data were generated in the Penn Cytomics and Cell Sorting Shared Resource Laboratory at the University of Pennsylvania (RRID:SCR_022376). Penn Cytomics is partially supported by the Abramson Cancer Center NCI Grant (P30 016520). Multiple figures were created using Biorender.com under license.

We thank the PENN-CHOP Human Lung Tissue Bank Pipeline (Maria Basil, M.D, Ph.D.; Edward Morrisey, Ph.D.) for the provision of human lung tissue sections.

