## Supplemental Data for "REGULATORY T CELLS PROTECT AGAINST ABERRANT REMODELING IN A MOUSE MODEL OF PULMONARY FIBROSIS"

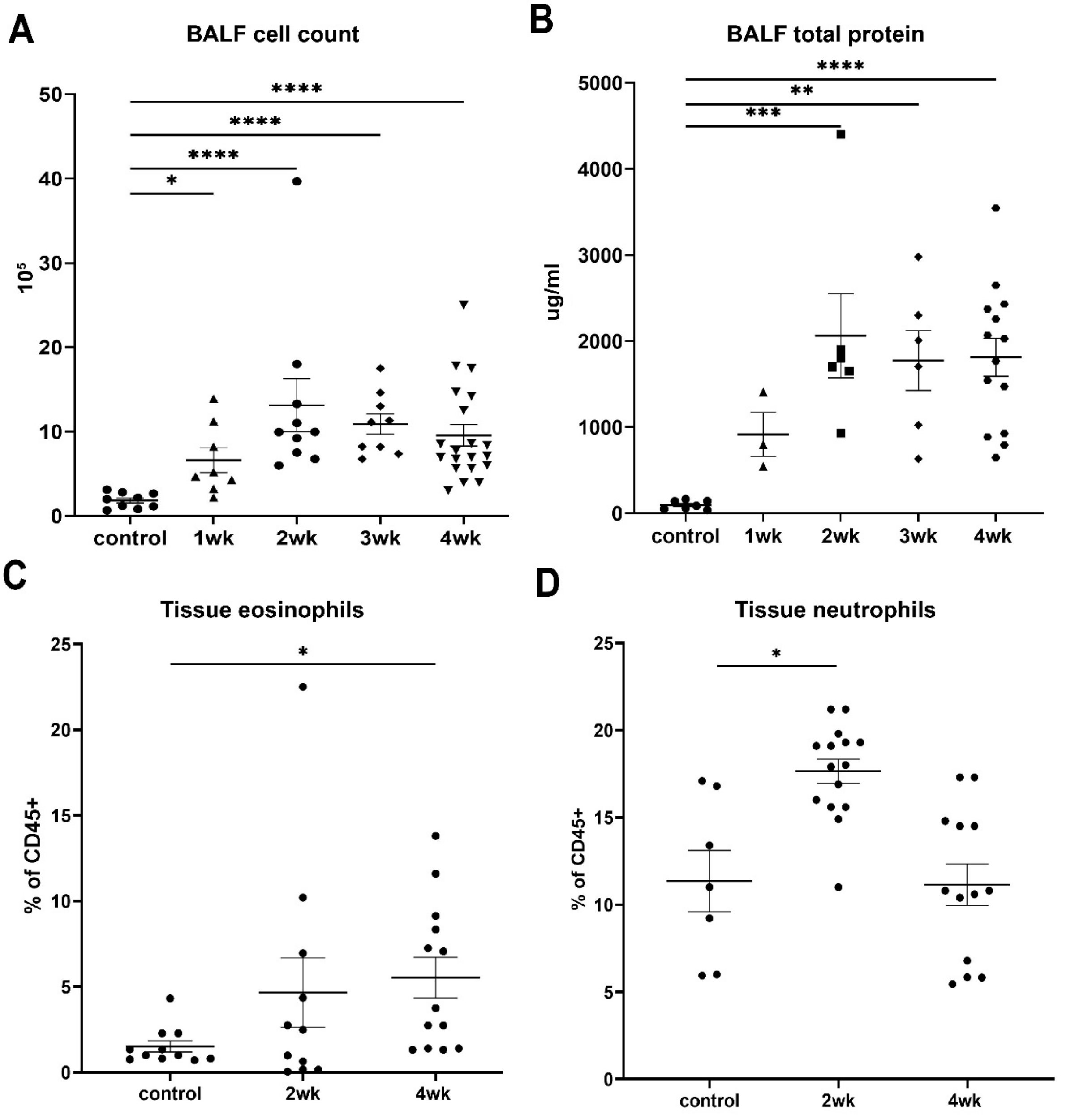

**Supplemental figure 1: AT2-specific lung injury induces alveolitis** Bronchoalveolar lavage fluid (BALF) collected at defined time points after two oral doses of tamoxifen in SP-$C^{I73T}$ and control mice showed increased (A) total cell counts and (B) protein concentrations. Flow cytometric analysis of lung tissue at 2- and 4-weeks post-injury revealed elevated myeloid cell populations, including (C) eosinophils and (D) neutrophils. Statistical analysis was performed using one-way ANOVA (*$p < 0.05$, **$p < 0.005$, ***$p = 0.0009$, ****$p < 0.0001$). SP-$C^{I73T}$, n = 3–20; control, n = 7–11.

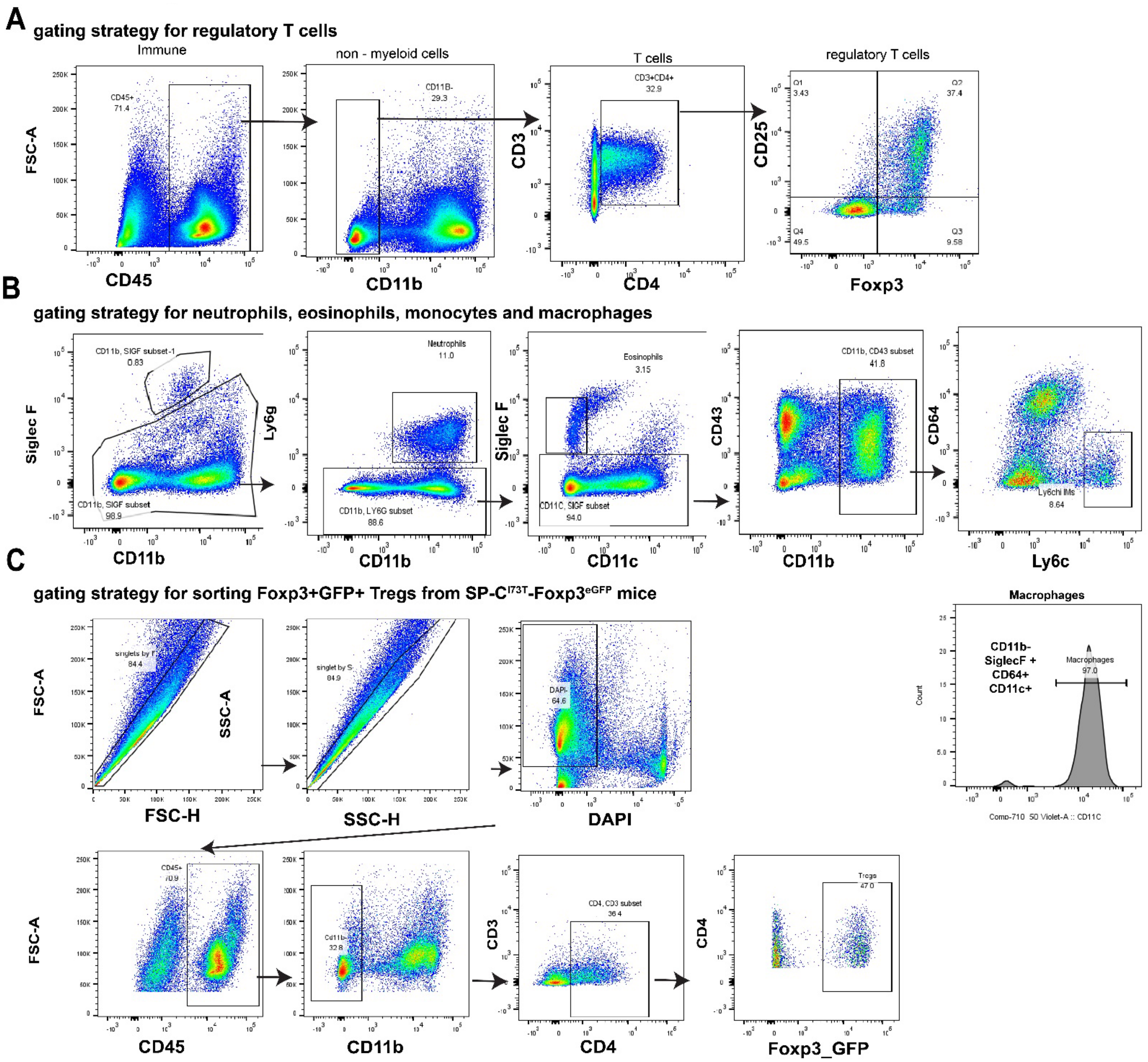
A
gating strategy for regulatory T cells
Immune
non - myeloid cells
T cells
regulatory T cells
FSC-A
CD45
CD45+
71.4
CD11b
CD11B-
29.3
CD3
CD4
CD3+CD4+
32.9
CD25
Foxp3
Q1
3.43
Q2
37.4
Q3
9.58
Q4
49.5
B
gating strategy for neutrophils, eosinophils, monocytes and macrophages
Siglec F
CD11b, SIGF subset-1
0.83
CD11b, SIGF subset
98.9
Ly6g
Neutrophils
11.0
CD11b, LY6G subset
88.6
Eosinophils
3.15
CD11C, SIGF subset
94.0
CD11c
CD43
CD11b, CD43 subset
41.8
CD64
Ly6c
Ly6chi IMs
8.64
C
gating strategy for sorting Foxp3+GFP+ Tregs from SP-C$^{I73T}$-Foxp3$^{eGFP}$ mice
singlets by F
84.4
FSC-H
SSC-A
singlet by S
84.9
SSC-H
DAPI-
64.6
DAPI
Macrophages
CD11b-
SiglecF +
CD64+
CD11c+
Macrophages
97.0
Count
Comp-710_50 Violet-A :: CD11C
CD45+
70.9
Cd11b-
32.8
CD4, CD3 subset
36.4
Tregs
47.0
Foxp3_GFP

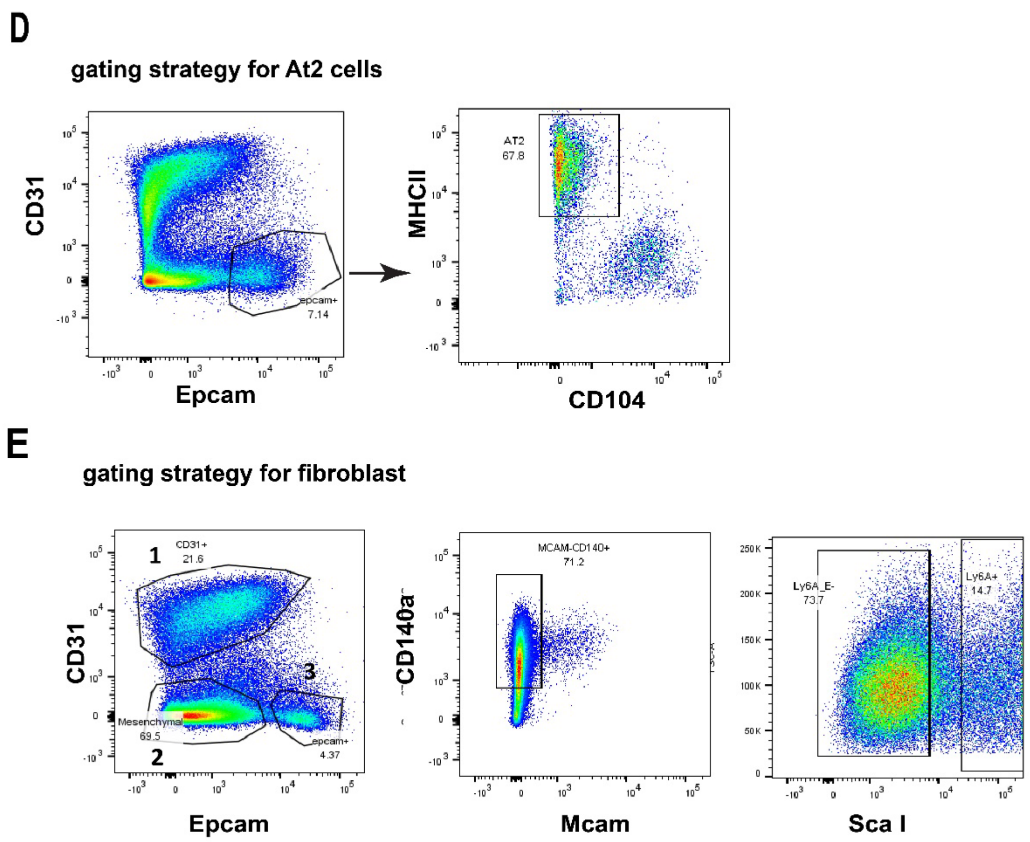

**Supplemental figure 2: Gating strategy for lung immune, epithelial, and mesenchymal cell populations.** Flow cytometric gating and phenotyping were used to identify (A) regulatory T cells, (B) neutrophils, eosinophils, monocytes, and macrophages in SP-C$^{I73T}$ mouse lungs, and (C) regulatory T cells in SP-C$^{I73T}$Foxp3$^{eGFP}$ mice. Epithelial and mesenchymal populations included (D) alveolar type 2 (AT2) cells and (E) adventitial fibroblasts.

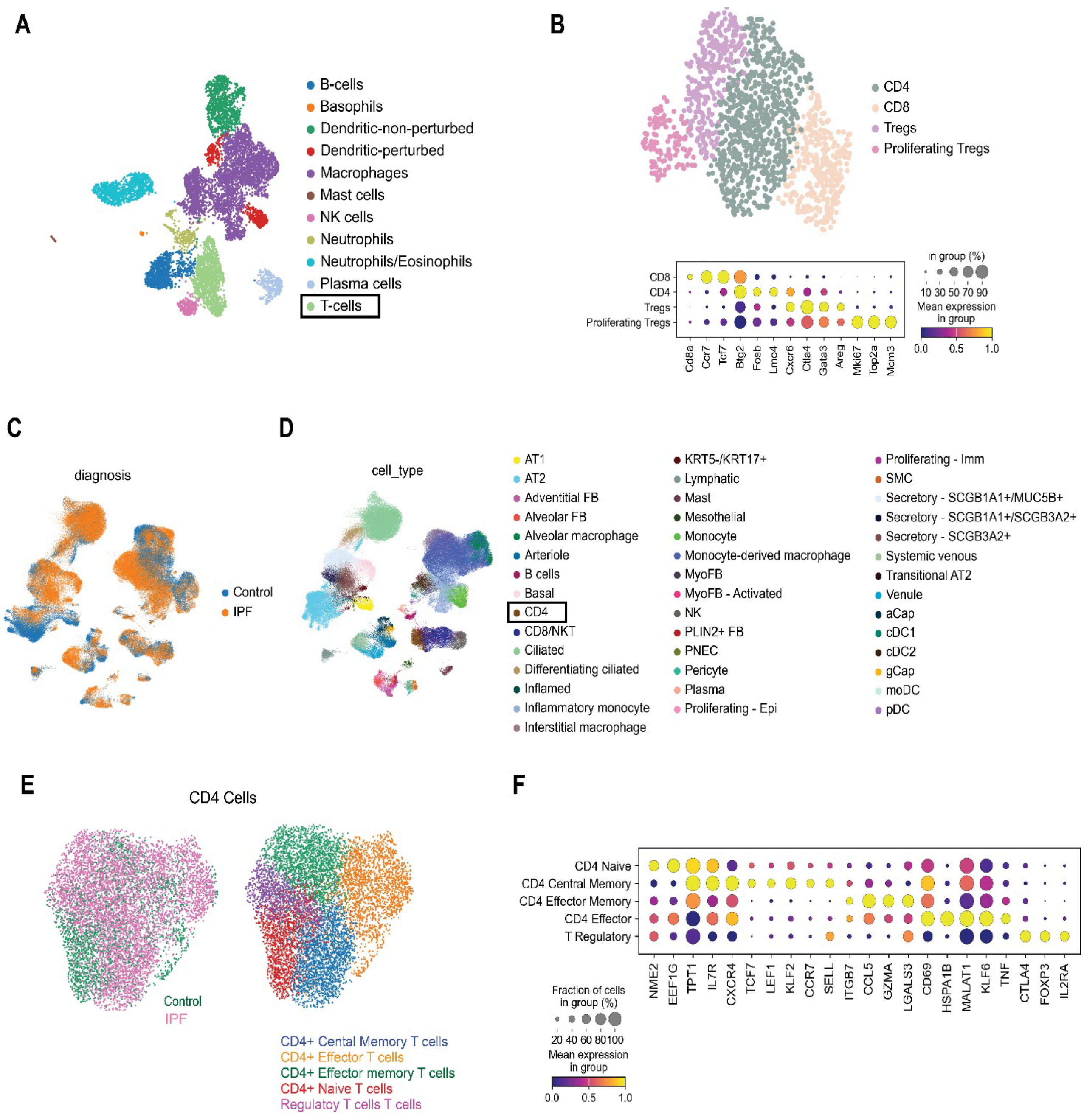

**Supplemental Figure 3:** Marker Genes Used to Identify Tregs in Murine and Human scRNA Data Sets A) Annotated UMAP of CD45 positive cells in previously published GSE234604 identifying major immune populations in *Sftpc*$^{I73T}$ mouse lungs. B) Re-clustered UMAP of T-cells and gradient dot plot with marker genes used to define each T-cell subcluster. C) UMAP of IPF and control lungs from previously published human lung data set GSE227136 annotated by disease status D) Cell type annotation of GSE227136 as published by authors and highlighting a subset of cells that are mapped as CD4 T-cells. E) UMAP of CD4 T-cells in GSE227136 denoting disease status and newly annotated subclusters of CD4 T-cells. F) Gradient dot plot of marker genes used to identify and annotate the subclusters of CD4 T-cells.

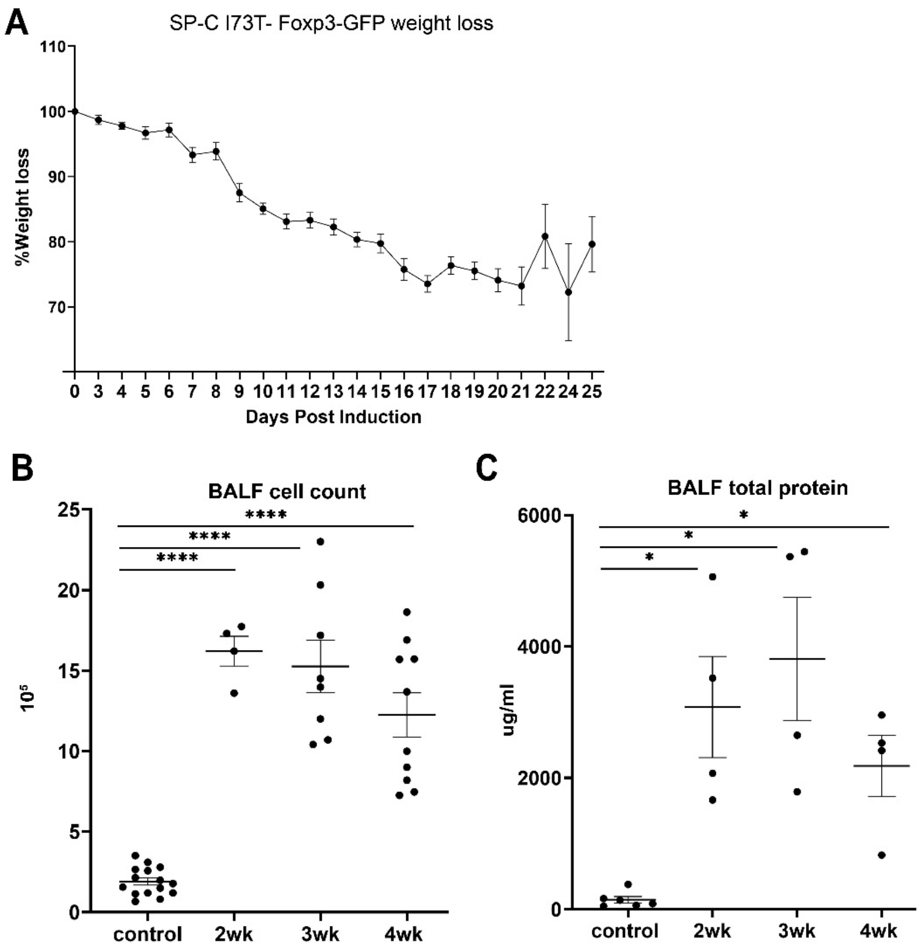

**Supplemental figure 4: Generation of the SP-C$^{I73T}$Foxp3$^{eGFP}$ mouse model to recapitulate AT2-specific lung injury.** Tamoxifen administration in SP-C$^{I73T}$Foxp3$^{eGFP}$ mice induced (A) progressive weight loss following AT2-specific mutant SP-C expression. Increased (B) total BAL cell counts and (C) protein concentrations at indicated time points indicate alveolitis and fibrosis, consistent with the phenotype observed in SP-C$^{I73T}$ mice. Statistical analysis was performed using one-way ANOVA (*$p < 0.05$, ****$p < 0.0001$). SP-C$^{I73T}$, n = 4–10; control, n = 6–14**.**

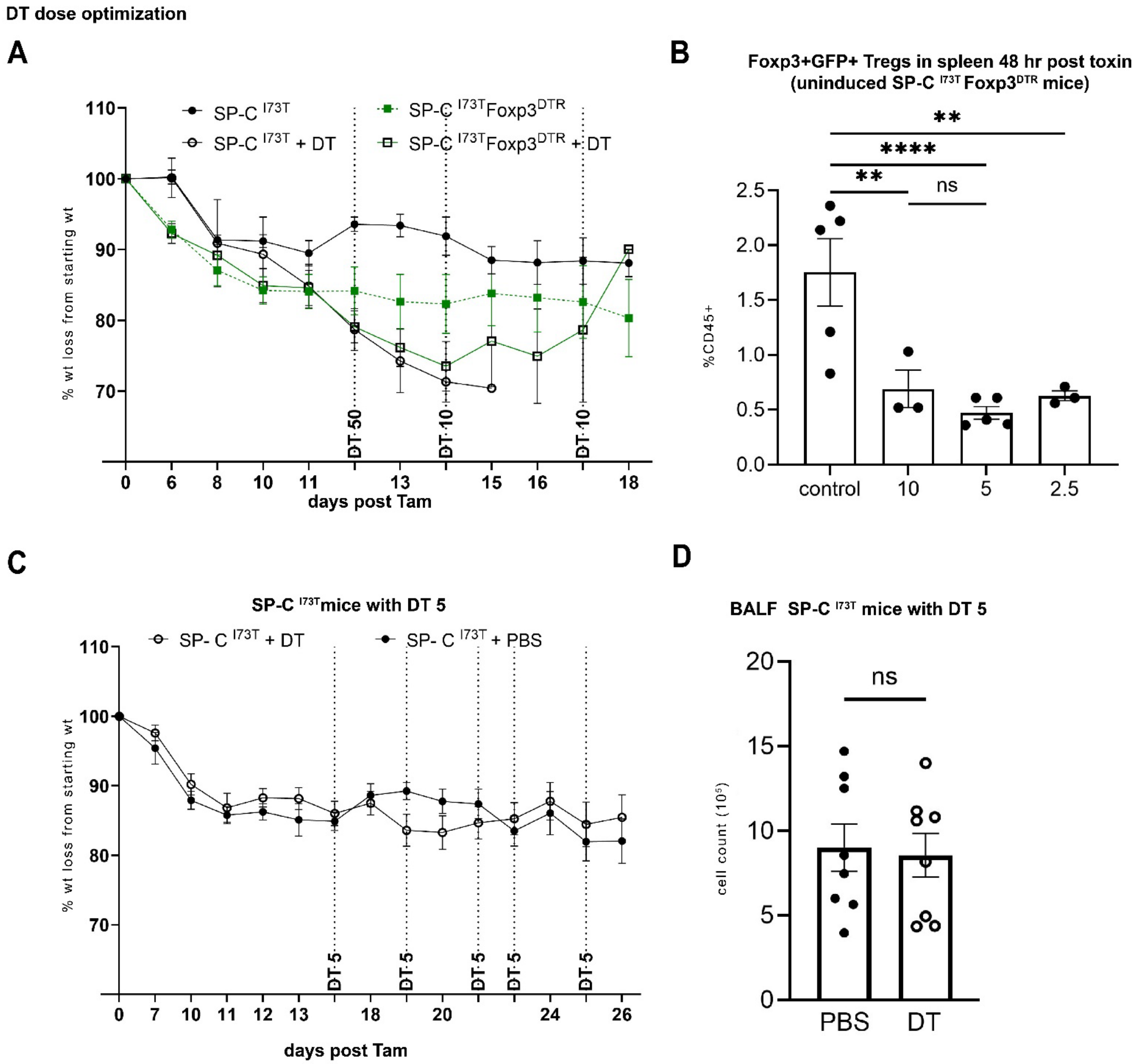

**Supplemental figure 5: Diphtheria toxin dose (DT) optimization and toxicity studies in SP-C$^{I73T}$ and SP-C$^{I73T}$-Foxp3$^{DTR}$ mice** (A) SP-C$^{I73T}$-Foxp3$^{DTR}$ and SP-C$^{I73T}$ mice received intraperitoneal DT injections (50, 10, and 10 μg/kg) beginning 12 days post-tamoxifen induction, with smaller doses administered every other day. Daily weight monitoring showed >30% weight loss in both groups. (B) Flow cytometric analysis of splenic CD45+ immune cells 48 hours post-DT administration (10, 5, or 2.5 μg/kg) assessed Treg depletion efficiency in SP-C$^{I73T}$-Foxp3$^{DTR}$ mice. (C) Weight loss and (D) BALF total cell counts were measured in SP-C$^{I73T}$ mice treated with either 5 μg/kg DT every other day or PBS. Statistical analysis was performed using one-way ANOVA (**p =0.001, ****p < 0.0001). SP-C$^{I73T}$Foxp3$^{DTR}$ + DT n=7; SP-C$^{I73T}$ + DT n=6; SP-C$^{I73T}$Foxp3$^{DTR}$ + PBS n=4; SP-C$^{I73T}$+ PBS n =4.

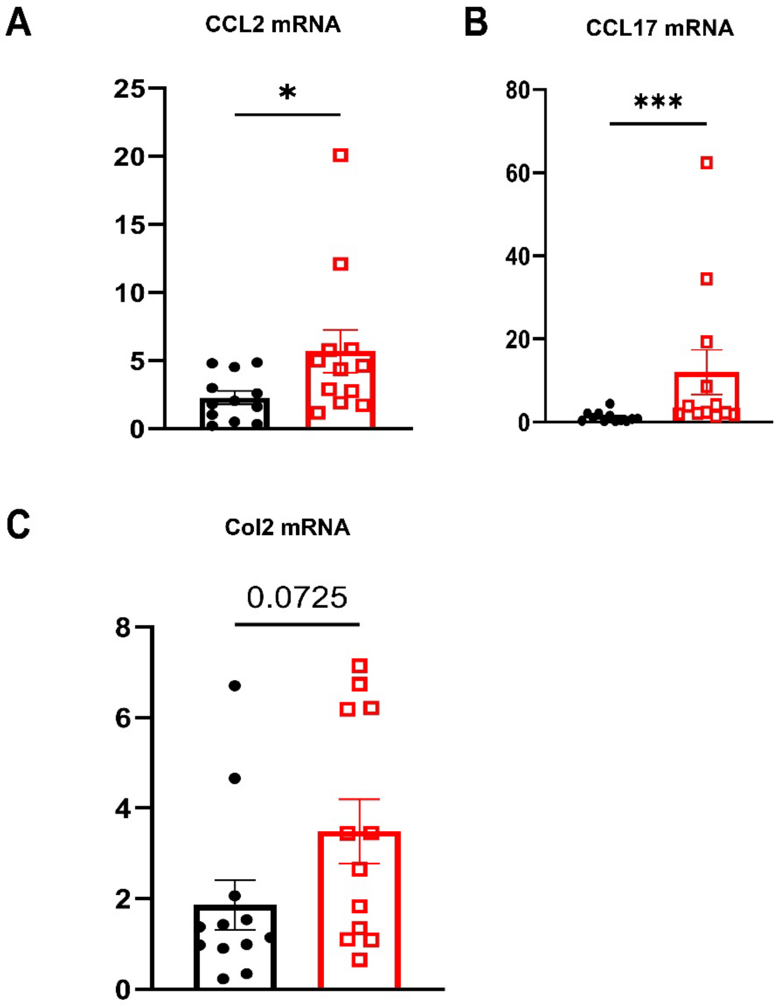

**Supplemental figure 6: Treg depletion during the fibrogenic phase exacerbates disease in SP-C$^{I73T}$Foxp3$^{DTR}$ mice.** Mice received intraperitoneal injections of diphtheria toxin (5 µg) or PBS every 48 hours, starting on day 17 post-tamoxifen induction. On day 26, whole lung tissue was collected from DT-treated (n = 9) and PBS-treated (n = 9) SP-C$^{I73T}$Foxp3$^{DTR}$ mice for gene expression analysis by qRT-PCR, pro-inflammatory chemokines (A) *Ccl2* and (B) *Ccl17*, which mediate monocyte and macrophage recruitment, as well as (C) the collagen-associated gene *Col2a1*, were elevated in DT-treated mice. Statistical analysis was performed using Unpaired t-test *p<0.05 *** p<0.001.

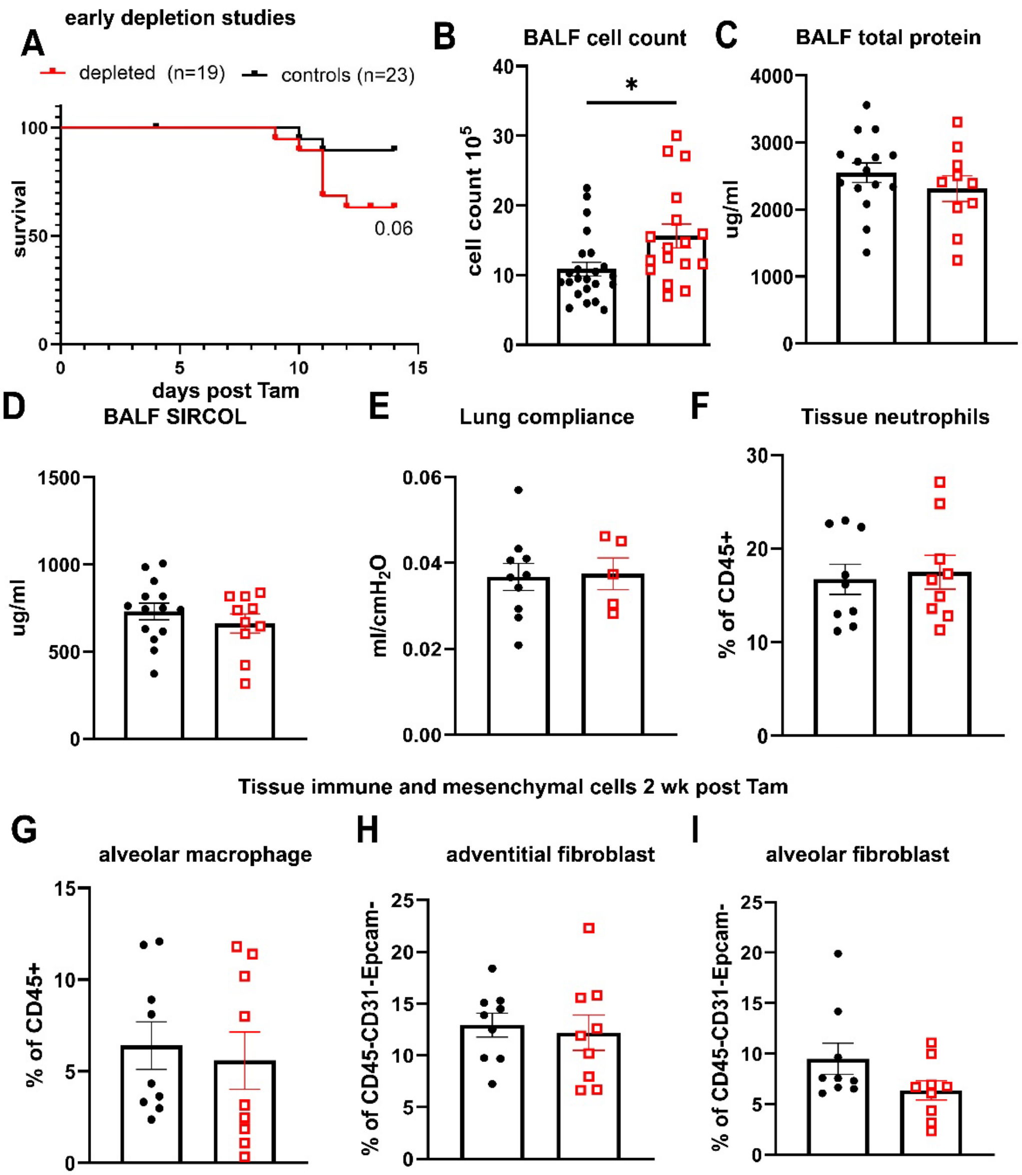

**Supplemental Figure 7: Early Treg depletion does not alter fibrotic endpoints in SP-C$^{I73T}$Foxp3$^{DTR}$ mice following tamoxifen induction.** Mice received intraperitoneal injections of diphtheria toxin (DT; 5 µg) or PBS every 48 hours, starting 2 days prior to tamoxifen (Tam) induction (A) Kaplan–Meier survival curves of SP-C$^{I73T}$Foxp3$^{DTR}$ mice treated with PBS (n=23) or DT (n=19); log-rank (Mantel–Cox) test, $p$ = 0.0648. (B–C) Total bronchoalveolar lavage fluid (BALF) cell counts and protein levels at day 15 post-Tam induction (PBS: n=23; DT: n=19); unpaired $t$-test with Welch's correction, $p$ = 0.01. (D-E) fibrotic endpoints at day 15: (D) BALF soluble collagen measured by Sircol assay (PBS: n=14; DT: n=10); (E) static lung compliance (Cst) (PBS: n=10; DT: n=5). (F–I) Flow cytometric analysis of lung immune (neutrophils, alveolar macrophages) and mesenchymal (adventitial and alveolar fibroblasts) populations on day 15 post-Tam (PBS: n=9; DT: n=9).

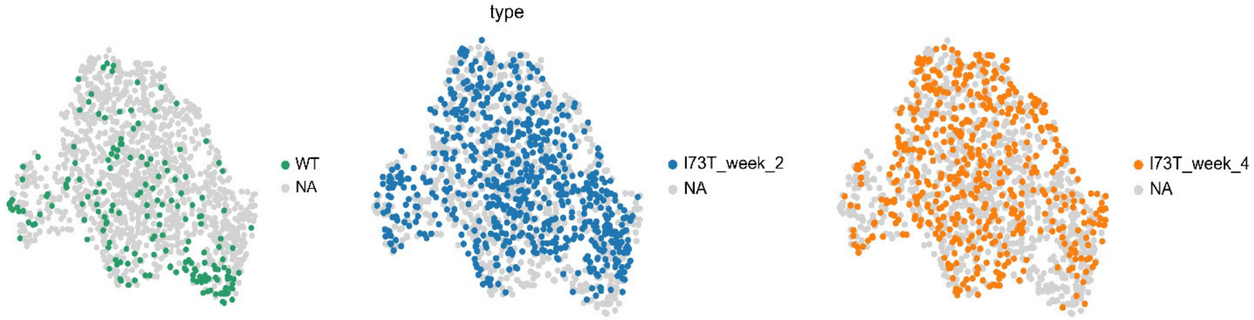

**Supplemental Figure 8: Increased expression of T cell–specific genes in SP-C$^{I73T}$ mice**. Single-cell RNA sequencing (scRNA-seq) was performed on lungs from control and SP-C$^{I73T}$ mice at 2- and 4-weeks post-tamoxifen induction (GSE234604), as described in Methods. (A) UMAP analysis of 1,341 CD3+ T cells reveals increased expression of T cell–specific genes following mutant SP-C expression. *n* = 2 mice per genotype per time point**.**

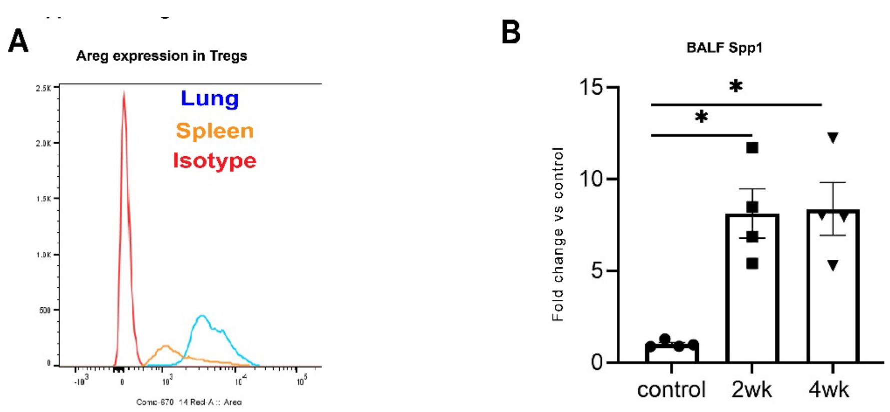

**Supplemental Figure 9: Lung Tregs produce pro-repair growth factors in SP-C$^{I73T}$ mice.** (A) Foxp3-GFP+ Tregs were FACS-purified from lungs and spleens of SP-C$^{I73T}$Foxp3$^{eGFP}$ mice 4 weeks post-tamoxifen induction, stimulated overnight with a cell stimulation cocktail (see Methods), and stained for intracellular Amphiregulin (Areg). Histograms show increased Areg expression only in lung-derived GFP+ Tregs following mutant SP-C expression. (B) BALF samples from control and SP-C$^{I73T}$ mice collected at 2- and 4-weeks post-induction were analyzed for Osteopontin (Spp1), a protein involved in growth, repair, and IPF pathogenesis.

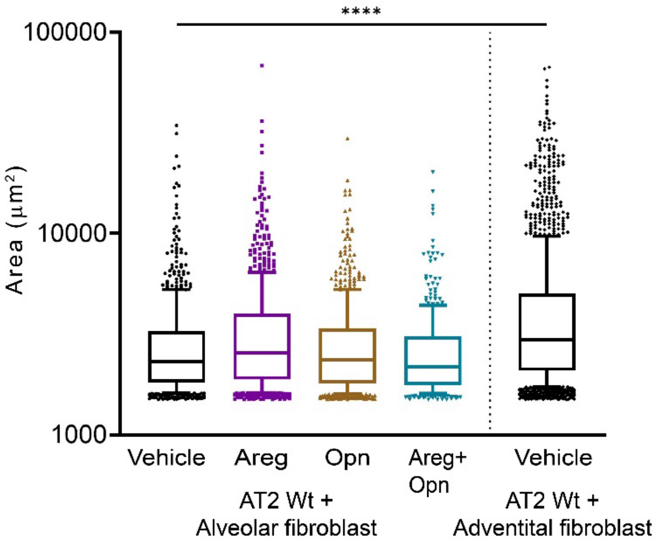

**Supplemental Figure 10: Lung SP-C$^{I73T}$ T regs selectively enhance adventitial, but not alveolar, fibroblast-supported alveolar organoid growth in vitro post injury** Bi-cellular organoids containing FACS purified TdTom+ AT2 and ScaI+ adventitial fibroblast were cultured with either 100 nM Areg, 200 nM Opn, or Areg + Opn for 2 weeks. For comparison, tricellular organoids containing lung Tregs were prepared and co-cultured for 2 wks. Live cultures were imaged using EVOS FL Auto and images were quantified using ImageJ an “analyze particle” macro with minimum area threshold set to1500 $\mu m^2$. The experiment was independently repeated at least three times, each with three biological and three technical replicates ****$p<0.00005$ done by Ordinary one-way ANOVA.

| Supplemental Table 1: Antibodies for Flow Cytometry | | | | |
|---|---|---|---|---|
| Antibody | Fluor | Clone | Catalog Number | Manufacturer |
| CD45 | BB515 | 30F-11 | 564590 | Biolegend |
| CD45 | APC | 30F-11 | 103112 | Biolegend |
| CD25 | APC | PC61 | 561048 | BD Biosciences |
| CD25 | BV605 | PC61 | 563061 | BD Biosciences |
| Foxp3 | PE | FJK-16s | 12-5773-82 | Thermofisher |
| Foxp3 | APC | FJK-16s | 12-5773-82 | Thermofisher |
| CD3e | APC/fire 750 | 145-2C11 | 100361 | Biolegend |
| CD3e | PE/Cy7 | 145-2C11 | 561100 | BD Biosciences |
| CD4 | BV786 | GK1.5 | 5633331 | BD Biosciences |
| GATA3 | PE | TWAJ | 12-9966-42 | Invitrogen |
| Tbet | BV421 | 4B10 | 644815 | BD Biosciences |
| Rorgt | BV21 | Q31-378 | 562894 | BD Biosciences |
| CD62L | PE/Cy7 | MEL-14 | 104417 | BD Biosciences |
| Tigit | PE/Cy7 | 1G9 | 142107 | Biolegend |
| PD1 | BV711 | J43 | 744547 | Biolegend |
| GITR | PE/Cy7 | YGITR 765 | 120222 | Biolegend |
| CD103 | Percp cy5.5 | 2 E 7 | 121415 | Biolegend |
| CD44 | PE | IM7 | 553133 | BD Biosciences |
| Epcam | BV711 | G8.8 | 118233 | Biolegend |
| Epcam | BV785 | G8.8 | 118245 | BD Biosciences |
| CD31 | APC/Cy7 | MEC13.3 | 102533 | Biolegend |
| CD104 | PE/Cy7 | 346-11A | 123615 | Biolegend |
| CD51 | PE | RMV-7 | 104105 | Biolegend |
| SiglecF | PEGF594 | E50-2440 | 562757 | BD Biosciences |
| CD11b | BV421 | M1/70 | 101205 | eBiosciences |
| CD11c | BV711 | HL3 | 561022 | Biolegend |
| Ly6g | AF700 | 1A8 | 561236 | Biolegend |
| CD64 | PE/Cy7 | X54-5/7.1 | 139305 | Biolegend |
| CD43 | PE | S11 | 143205 | Biolegend |
| Ly6C | BV510 | HK1.4 | 128033 | Biolegend |
| CD3e | BUV395 | 145-2C11 | 563565 | BD Biosciences |
| Amphiregulin | | Poly goat IgG | BAF989 | R&D Systems |
| Biotin anti mouse Epcam | | G8.8 | 118204 | Biolegend |
| Biotin anti mouse CD140a | | APA5 | 135910 | Biolegend |
| Biotin anti mouse CD45 | | 30F-11 | 103104 | Biolegend |
| Biotin anti mouse CD31 | | MEC13.3 | 102504 | Biolegend |
| Anti mouse CD3 | | 17A2 | 16-0032-85 | Thermofisher |
| Anti mouse CD28 | | 37.51 | 16-0281-85 | Thermofisher |
| live/dead | BUV395 | viability dye | 423107 | Biolegend |
| live/dead | ef780 | viability dye | 65-0865-14 | Thermofisher |

| Supplemental Table 2: Primer Sequences | | |
|---|---|---|
| **Target** | **Forward** | **Reverse** |
| 18S | CGGCTACCACATCCAAGGAA | GCTGGAATTACCGCGGCT |
| b-actin | GGCACCACACCTTCTACAATG | GGGGTGTTGAAGGTCTCAAC |
| Col3a | CTAAAGGGGAGATGGCCTG | ATTATCGGGTTGGAGCGCAG |
| Col1a | GCAAGAGGCGAGAGAGGTTT | CAGTTCACACTCGTAGCCGT |
| Fgf7 | CATGCTTCCACCTCGTCTGT | CAGTTCACACTCGTAGCCGT |
| mKi67 | ACCATCATTGGACCGCTCCTT | GCCCTGATGAGTCTTGGCTA |
| Top2a | TACAGTGCTCAACCTCTGACG | GGGATCTCGTGTTGGGAAGG |
| Areg | GGTCTTAGGCTCAGGCCATTA | CGCTTATGGTGGAAACCTCTC |
| Ccl2 | Mm00441242_m1 | |
| Ccl17 | Mm01244826_g1 | |

| Supplemental Table 3: Antibodies for IF | | | |
|---|---|---|---|
| **Antibody** | **Catalog Number** | **Manufacturer** | **Dilution** |
| Brilliant Violet 605 Streptavidin | 405229 | Biolegend | 1 in 200 |
| APC Streptavidin | 405207 | Biolegend | 1 in 200 |
| Dapi (4',6-Diamidino-2-Phenylindole, Dihydrochloride) | D1306 | Thermofisher | 1 in 1000 |
| CD39L1/ETPD2 | AF5797 | R&D Systems | 1 in 200 |
| RAGE | 175410 | R&D Systems | 1 in 200 |
| Foxp3 | ab215206 | abcam | 1 in 200 |
| Foxp3 | 14-5773-82 | invitrogen | 1 in 200 |
| Npro2 | | In-house (refer to M&M) | 1 in 200 |
| Hoechst 33342 | H3570 | Invitrogen | 1 in 1000 |
| Goat anti Mouse - HRP | 170-6516 | BioRad | 1 in 10000 |
| Goat anti Rabbit - HRP | 170-6515 | BioRad | 1 in 10000 |
| AlexaFluor 647 Goat anti Rabbit | A32733TR | Invitrogen | 1 in 400 |
| AlexaFluor 488 Goat anti Rabbit | A32731 | Invitrogen | 1 in 400 |
| AlexaFluor 488 Goat anti Rat | A11006 | Invitrogen | 1 in 400 |
| AlexaFluor 674 Goat anti Rat | A21247 | Invitrogen | 1 in 400 |
